# Transcriptional responses to chronic oxidative stress require cholinergic activation of G-protein-coupled receptor signaling

**DOI:** 10.1101/2025.01.06.628021

**Authors:** Kasturi Biswas, Caroline Moore, Hannah Rogers, Khursheed A Wani, Arjamand Mushtaq, Read Pukkila-Worley, Daniel P Higgins, Amy K Walker, Gregory P. Mullen, James B Rand, Michael M Francis

## Abstract

Organisms have evolved protective strategies that are geared toward limiting cellular damage and enhancing organismal survival in the face of environmental stresses, but how these protective mechanisms are coordinated remains unclear. Here, we define a requirement for neural activity in mobilizing the antioxidant defenses of the nematode *Caenorhabditis elegans* both during chronic oxidative stress and prior to its onset. We show that acetylcholine-deficient mutants are particularly vulnerable to chronic oxidative stress. We find that extended oxidative stress mobilizes a broad transcriptional response which is strongly dependent on both cholinergic signaling and activation of the muscarinic G-protein acetylcholine coupled receptor (mAChR) GAR-3. Gene enrichment analysis revealed a lack of upregulation of proteasomal proteolysis machinery in both cholinergic-deficient and *gar-3* mAChR mutants, suggesting that muscarinic activation is critical for stress-responsive upregulation of protein degradation pathways. Further, we find that GAR-3 overexpression in cholinergic motor neurons prolongs survival during chronic oxidative stress. Our studies demonstrate neuronal modulation of antioxidant defenses through cholinergic activation of G protein-coupled receptor signaling pathways, defining new potential links between cholinergic signaling, oxidative damage, and neurodegenerative disease.

## INTRODUCTION

Organisms encounter a wide variety of deleterious internal and external stressors throughout their lifespan. The ability to effectively respond to these stress conditions is crucial for limiting damage and enhancing organismal survival. Transcriptional activation or repression of stress-responsive gene networks in specific cells or tissues is a key component of organismal protective strategies. For instance, adaptive transcriptional responses are important for protection from pathogen infection or exposure to exogenous chemicals and toxins (Cao, 2016; Hu et al., 2023; Irazoqui et al., 2010; Nandi and Aroeti, 2023). Moreover, impaired stress responses contribute to human age-related diseases, including neurodegenerative disorders such as Alzheimer’s disease (Biswas et al., 2022; Ma et al., 2011; Manoharan et al., 2016).

Defining gene regulatory networks that limit damage in response to stress and the mechanisms for their coordination across tissues is therefore an area of intense interest. The nervous system has recently emerged as an important nexus in coordinating systemic stress responses, such as the mitochondrial unfolded protein response (UPR^mt^) (Bar-Ziv et al., 2023; Liu et al., 2022; Wani et al., 2020). However, the extent of nervous system participation in coordinating responses to other classes of environmental stressors is not clearly defined.

Oxidative stress is among the most significant and damaging stresses that organisms must face. It occurs as a result of the aberrant accumulation of reactive oxygen species (ROS) and is an important aggravating factor in the pathogenesis of a variety of diseases, including cancer, diabetes, and neurodegenerative disease (Bhatti et al., 2022; Biswas et al., 2022; Valko et al., 2004; Zhang et al., 2011). During oxidative stress, excess ROS cause cellular damage by reacting with biomolecules such as proteins and lipids (Back et al., 2012; Juan et al., 2021; Paulsen et al., 2011; Yang et al., 2016). A major adverse consequence of both oxidative stress and neurodegenerative disease is an accumulation of insoluble toxic protein aggregates, which occur as a result of perturbed protein homeostasis (proteostasis) and produce further proteostasis disruptions (Hipp et al., 2019). To protect against oxidative stress, eukaryotes activate sophisticated transcriptional responses for the upregulation of defense systems that cope with elevated ROS levels and their deleterious consequences (Nguyen et al., 2009); however, our understanding of how response programs for protection from oxidative stress are coordinated across tissues remains quite limited.

Here, we investigate the role of neuronal signaling in mobilizing the transcriptional response to oxidative stress in the nematode *Caenorhabditis elegans. C. elegans* offers clear strengths for investigating oxidative stress responses and their regulation by the nervous system. Importantly, both neurotransmitter utilization (Bargmann, 1998) and the core stress response pathways are highly conserved from worms to humans (Blackwell et al., 2015; Murphy et al., 2003; van Heemst, 2010). Moreover, genetic disruptions in neural function are well-tolerated, allowing straightforward assessment of specific neuron classes, transmitter systems, and receptor types (Bargmann, 1998; Chun et al., 2015). Finally, survival assays for quantifying the organismal impacts of oxidative stress (Jia and Sieburth, 2021; Senchuk et al., 2017; Van Raamsdonk and Hekimi, 2009) and biochemical methods to assess oxidative damage (Goudeau and Aguilaniu, 2010) are well-established. By combining straightforward stress assays with transcriptome profiling and the disruption of specific neurotransmitter systems, we explore organismal-level links between neuronal function and conserved pathways for oxidative stress regulation.

We show that extended oxidative stress mobilizes a broad transcriptional response, including the upregulation of canonical antioxidant and detoxification pathways, transcription factors, and genes involved in protein homeostasis. The transcriptional response to oxidative stress, most notably upregulation of genes associated with protein degradation through the proteasome, is severely blunted by either a deficiency of cholinergic neurotransmission or by mutation of the Gq-coupled muscarinic acetylcholine receptor (mAChR) *gar-3* gene. Disruption of ACh signaling and mutation of *gar-3* produce similar decreases in survival during chronic oxidative stress, while specific overexpression of *gar-3* in cholinergic motor neurons offers protection. Together, our findings demonstrate the importance of neuronal cholinergic G_q_-coupled GPCR signaling for coordinating a cytoprotective oxidative stress response transcriptional program.

## RESULTS

### Neuronal silencing increases organismal vulnerability to chronic oxidative stress

To determine whether neurotransmission may impact organismal vulnerability to chronic oxidative stress, we measured the effects of neuronal silencing on longevity in the presence of the redox cycler paraquat. We first compared the longevity of wild-type animals treated with paraquat (PQ, 4 mM) from day 1 of adulthood. Consistent with prior work (Dues et al., 2016; Schaar et al., 2015; Senchuk et al., 2017; Van Raamsdonk and Hekimi, 2012), we found that wild-type animals live up to ∼25 days under normal conditions, compared with approximately 15 days in the continued presence of PQ (**Fig. S1A,B**). The decrease in survival due to PQ treatment was partially reversed by supplementation with N-acetyl cysteine (NAC), a chemical precursor to the antioxidant glutathione (Desjardins et al., 2017) (**Fig. S1C**). To further implicate oxidative stress in the reduced longevity observed with PQ treatment, we next measured the survival of mutants deficient for the antioxidant gene superoxide dismutase 2 (*sod-2*) which are particularly susceptible to oxidative damage (Van Raamsdonk and Hekimi, 2009). The survival of *sod-2* mutants in the presence of PQ was significantly decreased compared to wild type controls (by nearly 50%) (**Fig. S1D)**. Finally, we quantified protein carbonylation following extended PQ exposure as a measure of oxidative damage. We found that a 48 hr exposure to 4 mM PQ produces a striking increase in carbonylated/oxidized proteins (**Fig. S1E**). Together, our data demonstrate that extended PQ exposure substantially increases oxidative stress and damage, leading to reduced organismal survival.

Using pan-neuronal expression of the histamine-gated chloride channel HisCl1 to silence neuronal activity (Pokala et al., 2014), we sought to investigate the contribution of neuronal signaling to survival under oxidative stress conditions. We first asked whether neuronal silencing during the early stages of oxidative stress exposure altered organismal survival. Specifically, we exposed 1-day adults to 4 mM PQ for 48 hr prior to neuronal silencing (24 hr) in the continued presence of PQ. We found that neuronal silencing significantly decreased the survival of PQ-treated animals compared to PQ-treated controls that did not undergo neuronal silencing (**Fig. 1A**). Importantly, the same period of silencing did not significantly affect lifespan in the absence of PQ as measured by IP50, though a small reduction was detected from analysis of the survival curve (**Fig. S2A**). These results demonstrate that neuronal silencing during the early stages of chronic oxidative stress increases organismal susceptibility, suggesting that neuronal activity may play a protective role during chronic oxidative stress.

**Figure 1.**
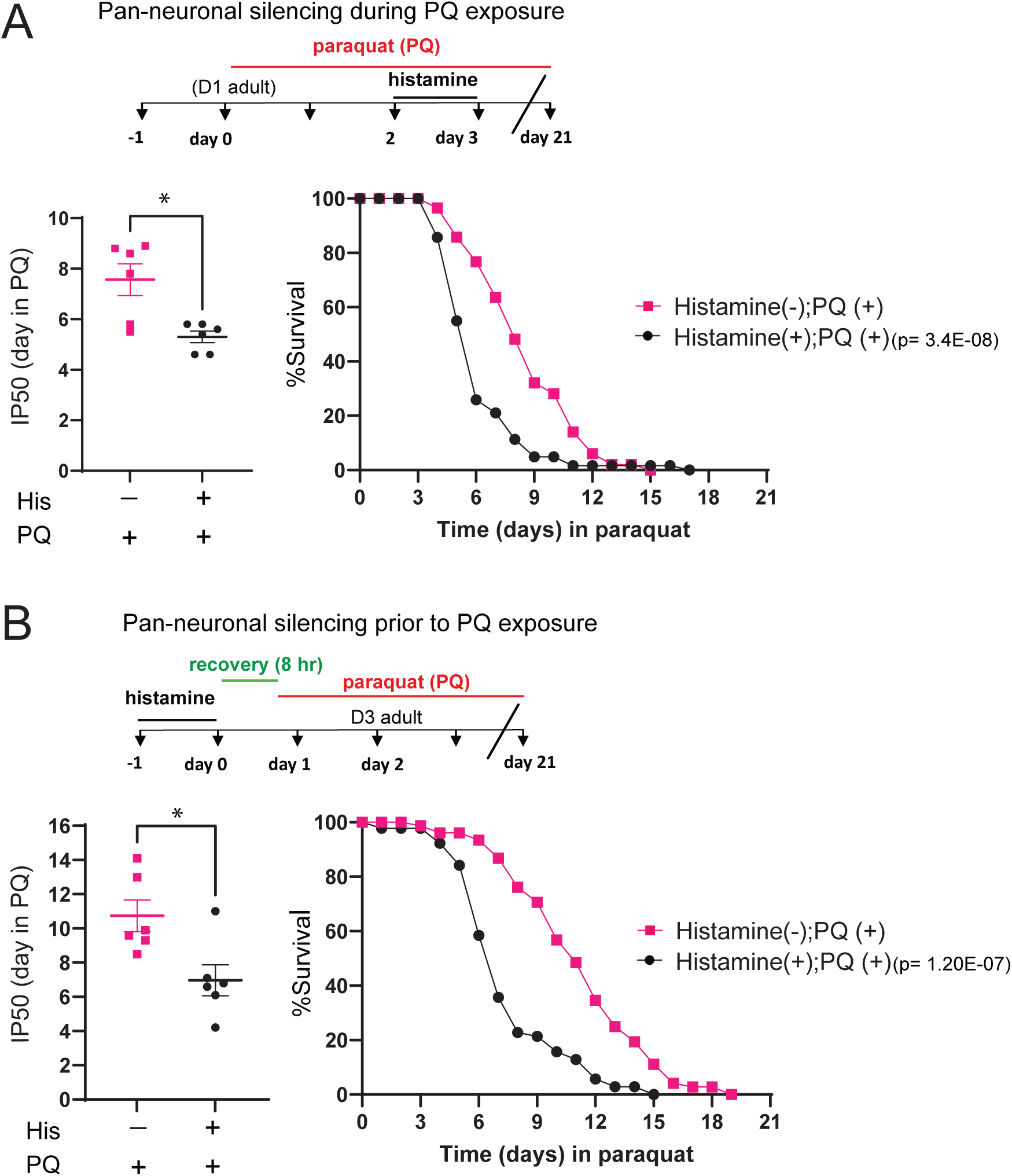
Neuronal activity protects from chronic oxidative stress **(A)** Top, experimental timeline indicating timing and duration of histamine and PQ treatment for survival assays. Left: Bar graph comparing IP50 measurements for control (His-) and His-treated (His+) *kyEx4571* (*tag-168p*::HisCl1::SL2::GFP) transgenic animals in the presence of PQ. The day at which His-treated animals reach half of their initial population is significantly advanced compared to control, indicating reduced survival. Each point represents one independent trial. Bars indicate mean ±SEM. *\*p=0.0135*, unpaired two-tailed t-test with Welch’s correction. Number of independent trials: n=6. Right: Kaplan-Meier survival curves comparing His+ and His- groups. Pan-neuronal silencing (24 hr) during oxidative stress reduces survival in the continued presence of PQ. Kaplan-Meier curves are cumulative of 6 trials. *\*\*\*\*p= 3.4E-08*, Fisher’s exact test at 50% of survival. Total number of animals: n= 59 (His-), 64 (His+). **(B)** Top, experimental timeline indicating timing and duration of histamine and PQ treatment for survival assays. Note 8 hr recovery period between the end of histamine treatment and the onset of PQ exposure. Left: Bar graph comparing (IP50 measurements for control (His-) and His-treated (His+) *kyEx4571* (*tag-168p*::HisCl1::SL2::GFP) transgenic animals in the presence of PQ. The day at which His-treated animals reach half of their initial population is significantly advanced compared to control, indicating reduced survival. Each point represents one independent trial. Bars indicate mean ± SEM. *\*p=0.0156*, unpaired two-tailed t-test with Welch’s correction. Number of independent trials: n=6. Right: Kaplan-Meier survival curves comparing His+ and His- groups. Pan-neuronal silencing (24 hr) prior to the onset of oxidative stress reduces organismal survival in the continued presence of PQ. Kaplan-Meier curves are cumulative of 6 trials. *\*\*\*\*p= 1.20E-07*, Fisher’s exact test at 50% of survival. Total number of animals, n= 82 (His-), 89 (His+).

To determine when neuronal activity may be most important, we next assessed the effects of neuronal silencing prior to PQ exposure. Specifically, we silenced neuronal activity for 24 hr followed by an 8 hr recovery period, prior to PQ exposure (**Fig. 1B**). Surprisingly, we found that neuronal silencing prior to the onset of oxidative stress also significantly decreased the survival of PQ-treated animals, while silencing prior to stress did not affect lifespan without PQ treatment, as measured by IP50 (**Fig. S2B**). Overall, our findings provide evidence that neural activity promotes organismal survival by protecting from oxidative damage. Interestingly, neuronal silencing prior to or during oxidative stress each reduce survival during chronic oxidative stress exposure, suggesting that activity may have roles both in acute responses to oxidative stress and in equipping the organism to mount an effective protective response.

### Disruption of either ACh or glutamate neurotransmission decreases survival to chronic oxidative stress

To determine whether specific neurotransmitter systems contribute preferentially toward this protective mechanism, we measured the survival of mutants deficient in various neurotransmitters, under chronic PQ exposure. We used available strains carrying mutations in biosynthetic machinery for specific neurotransmitters (dopamine, octopamine, serotonin, GABA, glutamate, acetylcholine). Critically, prior work demonstrated that the longevity of mutants deficient for any one of these signaling systems is not significantly reduced under control conditions, though disruption of GABA signaling produced a modest increase in lifespan (Chun et al., 2015). Of these, we found that disruption of either acetylcholine/ACh (VAChT/*unc-17* mutant) or glutamate (VGluT/*eat-4* mutant) transmission reduced survival during chronic oxidative stress most severely (**Fig. 2A**). Importantly, we also confirmed prior findings that mutation of either *unc-17* or *eat-4* did not significantly alter longevity under control conditions (**Fig. 2B**). Our results reveal the importance of signaling through the small molecule neurotransmitters ACh and glutamate in regulation of responses to chronic oxidative stress. In contrast, prior studies of acute oxidative stress implicated neuropeptide signaling in protection, while disruption of small molecule neurotransmitter signaling either had no appreciable involvement (Jia and Sieburth, 2021) or prolonged survival (Cornell et al., 2024). The differences in findings across acute and chronic stress may point towards intriguing differences in the way organisms respond to stress conditions of differing duration.

**Figure 2.**
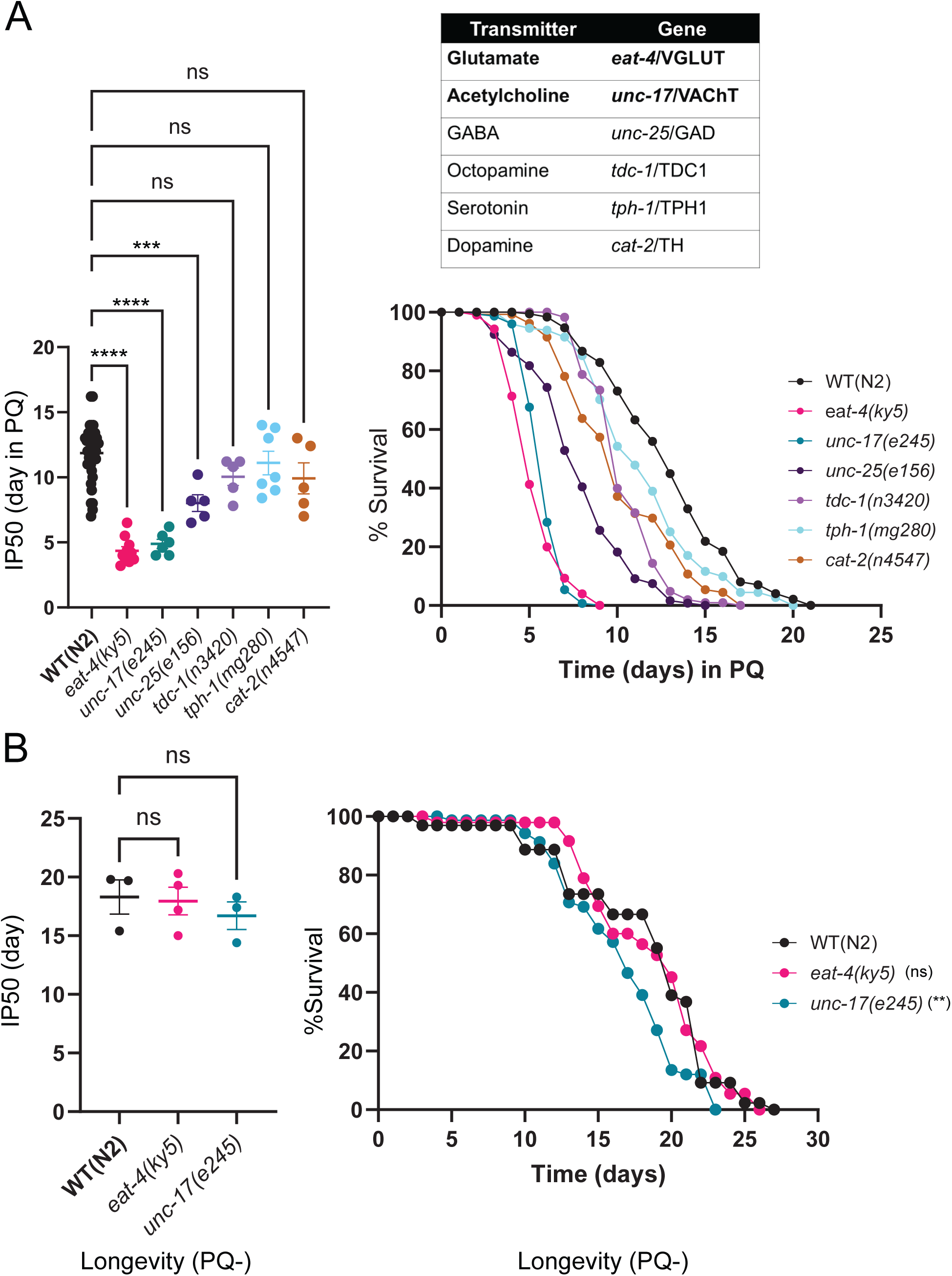
Disruption of either glutamate or acetylcholine neurotransmission increases vulnerability to PQ **(A)** Left: Bar graph comparing IP50 measurements for the indicated genotypes. The day at which *eat-4* (glutamate), *unc-17* (acetylcholine) or *unc-25* (GABA) mutants reach half of their initial population is significantly advanced compared to control, indicating reduced survival. The reduction in IP50 is greatest for *eat-4* and *unc-17* mutants. Each point represents one independent trial. Bars indicate mean ± SEM. *\*\*\*\*p <0.0001, eat-4(ky5); ****p<0.0001, unc-17(e245); ***p=0.0006, unc-25(e156); p=0.2718 (ns), tdc-1(n3420); p=0.9128 (ns), tph-1(mg280); p=0.2104 (ns), cat-2(n4547).* One-way ANOVA with Dunnett’s multiple comparisons test. Number of trials: n≥5. Right: Upper, table shows the six conserved neurotransmitters and corresponding mutant genes involved with either vesicular loading or neurotransmitter biosynthesis that were surveyed in the survival assays. Kaplan -Meier survival curves for wild type and indicated neurotransmitter-deficient mutants with chronic PQ exposure. Kaplan-Meier curves are cumulative of at least 5 trials. \*\*\*\**p=5.00E-06, eat-4(ky5); ****p=1.50E-12, unc-17(e245); ****p=2.40E-12, unc-25(e156); ****p=1.10E-08, tdc-1(n3420); p=0.2492 (ns), tph-1(mg280); ****p=4.30E-12, cat-2(n4547).* Fisher’s exact test at 75% of survival. Total number of animals: n= 527(N2), 251 (*eat-4*), 155 (*unc-17*), 148 (*unc-25*), 155 (*tdc-1*), 205 (*tph-1*), 145 (*cat-2*). **(B)** Left: IP50 comparisons showing glutamate- and acetylcholine-deficient mutants have no significant difference in their normal longevity. Each data point indicates an independent trial. Bars indicate mean ±SEM. *p =0.9712(ns), eat-4(ky5); p=0.6226 (ns), unc-17(e245).* One-way ANOVA with Dunnett’s multiple comparisons test. Number of trial ≥3. Right: Kaplan-Meier curve shows there is a modest reduction of longevity in *unc-17(e245)* and no difference in *eat-4(ky5)* compared to WT(N2). The difference in longevity for *unc-17(e245)* compared to WT(N2) was observed after 9 days. Kaplan-Meier curves are cumulative of at least 3 trials. *p =0.2425(ns), eat-4(ky5), **p=0.0012, (unc-17(e245)*. Fisher’s exact test at 50% of survival. Total number of animals: n=65, N2; 67, *eat-4(ky5)* and 80, *unc-17(e245)*.

As ACh is the most widely used small molecule neurotransmitter in *C. elegans*, we focused our subsequent analyses on the role of cholinergic signaling. Since a complete elimination of ACh function in *C. elegans* is lethal (Alfonso et al., 1993), we used the severe hypomorphic *unc-17(e245)* allele in our initial studies. *unc-17(e245)* animals are developmentally delayed, and the increase in vulnerability to PQ we observed may therefore have arisen indirectly from a more generalized requirement for cholinergic signaling in organismal health. To address this issue, we also tested animals carrying the *unc-17(e113)* allele that disrupts the COE motif in the *unc-17* promoter region (Kratsios et al., 2011; Serrano-Saiz et al., 2020) (J. Rand, unpublished). This mutation strongly decreases expression of *unc-17*/VAChT in cholinergic motor neurons that innervate body wall musculature but has less impact on *unc-17* expression in other cholinergic neurons and, therefore has more limited impacts on overall organismal development/health. *unc-17(e113)* animals displayed heightened vulnerability to PQ with a similar IP50 value to *unc-17(e245)* mutants (**Fig. 3A**). In contrast neither *unc-17* allele significantly reduced lifespan in the absence of PQ, as measured by IP50 (**Fig. 3B**). These results suggest that a loss of ACh release from cholinergic motor neurons may be sufficient to enhance vulnerability to oxidative stress. Consistent with this interpretation, specific knockdown of *unc-17* expression in cholinergic motor neurons by cell-specific RNAi also increased vulnerability to oxidative stress (**Fig. 3A**). Our results implicate ventral cord motor neurons as a key cellular source of ACh in mediating protection from oxidative damage.

**Figure 3.**
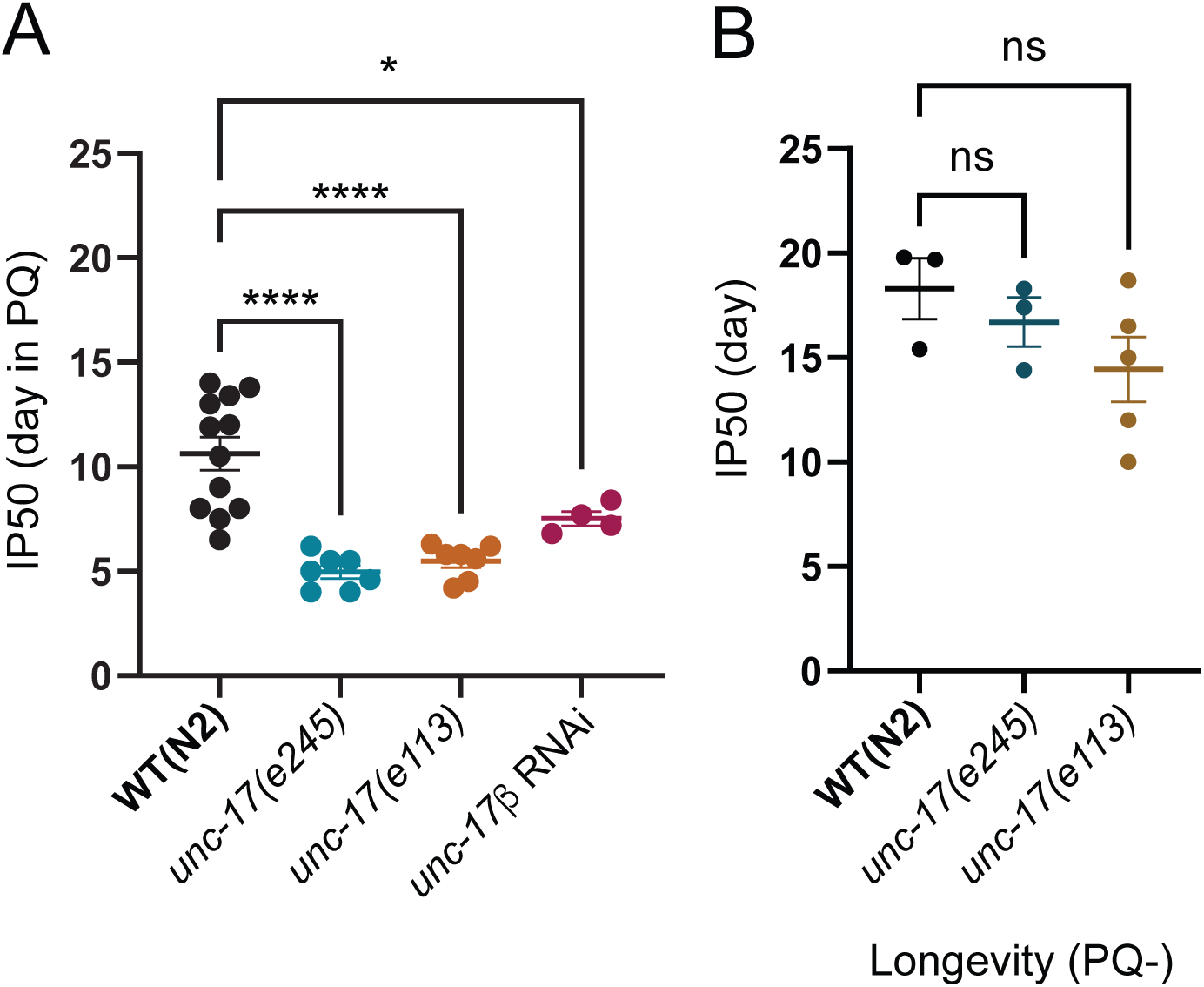
Disruption of cholinergic transmission from motor neurons reduces survival during chronic oxidative stress **(A)** Bar graph comparing IP50 measurements on PQ for wild type, *unc-17(e245)*, *unc-17(e113)*, or motor neuron-specific knockdown of *unc-17* by RNAi. *unc-17(e113)* mutants with reduced *unc-17* expression in motor neurons or RNAi downregulation of *unc-17* in motor neurons each show significant reductions in IP50, comparable to that of severely hypomorphic *unc-17(e245)* mutants. Each data point is one independent trial. Bars indicate mean ±SEM. *\*\*\*\*p <0.0001, unc-17(e245); ****p<0.0001, unc-17(e113); *p =0.0224*, *unc-17β RNAi*. One-way ANOVA with Dunnett’s multiple comparisons test. Number of trials, n≥4. Total number of animals: n= 303, N2; 200, *unc-17(e245);* 200, *unc-17(e113)*; and 91, *unc-17βp::unc-17* RNAi. **(B)** IP50 comparison of wild type (WT), *unc-17(e245),* and *unc-17(e113)* show there is no significant reduction in longevity under control (PQ-) conditions though *unc-17(e113)* is more variably affected. Each data point is one independent trial. Bars indicate mean ±SEM. *p= 0.7316 (ns), unc-17(e245); p=0.1826 (ns), unc-17(e113)*. One-way ANOVA with Dunnett’s multiple comparisons test. Number of trials: n≥3. Total number of animals: n= 65 WT(N2); 80, *unc-17(e245);* 63*, unc-17(e113)*.

### Extended oxidative stress mobilizes a broad transcriptional response

To better understand the impacts of chronic oxidative stress and the associated organismal responses, we next analyzed transcriptomic changes in response to extended PQ exposure (48 hr) by bulk RNA-seq. Specifically, we performed RNA sequencing of wild-type day 3 adults under control conditions, or following PQ exposure for 48 hrs. This timepoint is well before the occurrence of any lethality in our population survival assays. We found that extended oxidative stress remarkably affected the transcriptome (**Fig. 4A**). We observed broad expression changes (>2-fold, Padj< 0.01, FDR <0.01) encompassing more than 2,000 genes (1811 upregulated, 274 downregulated) after 2 days of PQ exposure (**Fig. 4B**, **Table S1**). To analyze the representation of functional gene categories amongst the PQ-upregulated genes, we compared up and downregulated genes using WormCat 2.0 (Holdorf et al., 2020) (**Fig. 4C**). Genes involved in stress or detoxification responses were highly enriched amongst the genes upregulated by PQ exposure and represent approximately 9% of the total gene set (Table S2). In particular, Phase 2 detoxification genes were highly represented among the upregulated genes, including the known SKN-1 target *gst-4* (Kahn et al., 2008; Tullet et al., 2008), ten other glutathione transferases (GSTs), and 22 UDP-glucuronosyl transferases (UGTs) (**Tables S1, S2**). Predicted Phase 1 detoxification genes, including 18 cytochrome p450 enzymes (CYPs) were also enriched amongst the upregulated genes. Finally, we note that the SKN-1-independent, hypoxia and pathogen-stress-responsive flavin-containing monooxygenase *fmo-2* gene is among the PQ-upregulated genes, providing evidence for the involvement of additional stress-responsive transcriptional pathways (Doering et al., 2022; Leiser et al., 2015; Wani et al., 2021).

**Figure 4.**
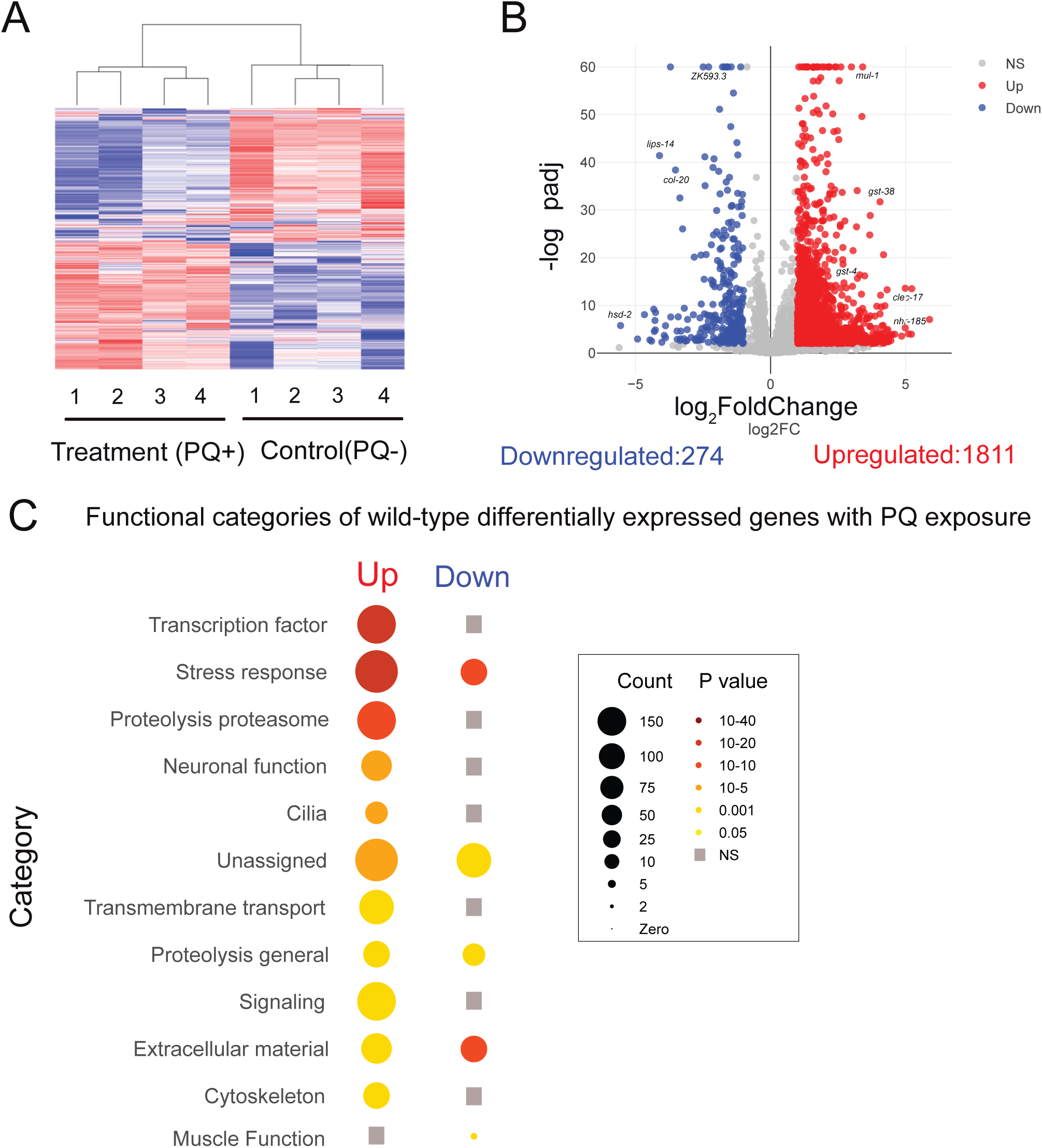
Extended oxidative stress mobilizes a broad transcriptional response (A) Heatmap shows clustering of treatment (PQ+) and control (PQ-) replicates for WT(N2). Number of replicates: n= 4 for both conditions. (B) Volcano plot (log_2_FoldChange, -log_10_padj) of differentially expressed genes following 48 hrs of PQ (4 mM) treatment compared to age-matched controls. Red: upregulated, Blue: Downregulated, Grey: not significantly different. Statistical cutoff for differential expression: FoldChange>2, P_adj_<0.01 and False Discovery Rate (FDR) <0.01. (C) Functional categorization of upregulated and downregulated genes that showed differential expression following 48 hr PQ exposure in wild type. The transcription factor, stress response and proteolysis proteasome categories were most strongly enriched amongst the differentially expressed genes. Grey squares indicate no significant enrichment in that category. Gene-count and P-value scales as indicated. A larger radius indicates a higher number of genes in that category. A darker color indicates a more significant P value.

The robust transcriptional response to extended PQ exposure showed considerable overlap with previously reported responses for acute exposure to other oxidants such as the metalloid arsenite (**Fig. 5A, Table S3)** (Oliveira et al., 2009), organic peroxide tBOOH (**Fig. 5B, Table S4**) (Oliveira et al., 2009), and the phenolic compound juglone (**Fig. 5C, Table S5**) (Wu et al., 2016), suggesting chronic and acute oxidative stress induce partially overlapping transcriptional responses. Arsenite mobilizes a largely SKN-1-dependent stress response, and tBOOH mobilizes a largely SKN-1-independent stress response (Goh et al., 2018; Oliveira et al., 2009). The overlap of both the arsenite and tBOOH transcriptomic responses with our findings suggests that both SKN-1-dependent and -independent pathways are mobilized in response to extended PQ oxidative stress.

**Figure 5.**
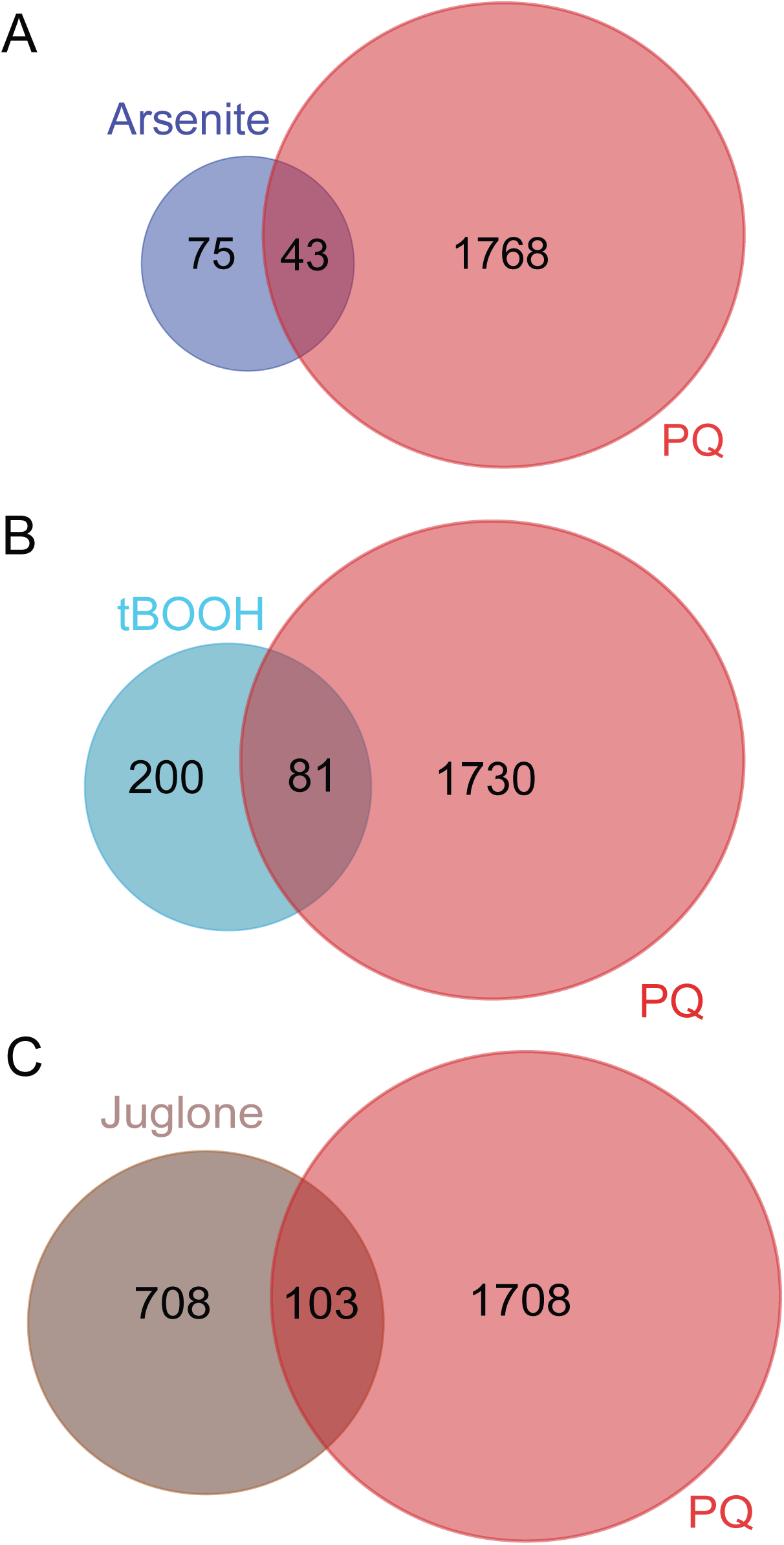
Comparison of transcriptomic responses between acute and extended oxidative stresses **(A)** Venn diagram shows the extent of overlap between genes upregulated in the presence of arsenite (Oliveira et al., 2009) compared with genes we identified as upregulated in response to extended PQ exposure. Approximately 36% (43 out of 118) of genes upregulated by arsenite treatment were also upregulated by PQ. Representative factor 3.5, p < 2.816e-14. **(B)** Venn diagram shows the extent of overlap between genes upregulated in the presence of tBOOH (Oliveira et al., 2009) compared with genes we identified as upregulated in response to extended PQ exposure. Approximately 29% (81 out of 281) of genes upregulated by tBOOH treatment are also upregulated by PQ. Representative factor 2.8, p<2.205e-18. **(C)** Venn diagram shows the extent of overlap between genes upregulated in the presence of juglone (Wu et al., 2016) compared with genes we identified as upregulated in response to extended PQ exposure. Approximately 13% (103 out of 811) of genes upregulated by juglone treatment are also upregulated by PQ. Representative factor 1.2, p<0.014.

The PQ-upregulated genes also included gene classes implicated in other stress-related processes. For instance, we identified numerous genes involved in proteolysis, including 120 predicted E3 ubiquitin ligases among the PQ-upregulated gene set. Additionally, many of the PQ-upregulated genes regulate transcription, including 50 predicted nuclear hormone receptor (*nhr*) genes and 38 genes encoding homeodomain transcription factors. As noted in prior oxidant response studies (Maremonti et al., 2019; Mesbahi et al., 2020; Sandhu et al., 2021), we found significant upregulation of genes encoding extracellular material, including collagen and cuticle proteins as well as CUB domain proteins. Interestingly, we also found significant enrichment of genes involved in neuronal function, including upregulation of genes involved in cholinergic signaling. In particular, we noted upregulation of the G-protein coupled muscarinic acetylcholine receptor (mAChR) *gar-3* gene (Hwang et al., 1999; Steger and Avery, 2004) and the *ric-3* gene that encodes a chaperone broadly important for the synthesis and synaptic delivery of ionotropic nicotinic acetylcholine receptors (nAChR) (Biala et al., 2009; Halevi et al., 2002). Curiously, among the 274 genes we identified to be downregulated by extended PQ exposure, we observed the strongest enrichment for collagen genes (24 genes) and other genes involved in detoxification (*gst-44*, *ugt-63*, *cyp-35B1*) and defense against pathogens (*thn-1*, *thn-2*, *irg-5*, C32H11.3). Overall, we conclude that the transcriptional response to extended PQ exposure induces many of the same gene classes previously implicated in both SKN-1-dependent and -independent responses to acute oxidant exposure. Additionally, we note that a more broad recruitment of stress-responsive pathways may arise due to distinct impacts of PQ as an oxidant, the extended duration of oxidant treatment used for our studies, or a combination of both factors.

### Cholinergic transmission is required for mobilization of the transcriptional response to extended oxidative stress

Encouraged by our results indicating increased vulnerability to oxidative stress in mutants where cholinergic neurotransmission is disrupted, we next sought to define the involvement of cholinergic signaling in the transcriptional response to extended PQ exposure. As described above for wild type, we performed RNA sequencing of day-3 adult *unc-17(e113)* mutants under control conditions or following 48 hr of 4 mM PQ exposure. We found that extended oxidative stress mobilized a significant transcriptional response in *unc-17* mutants (**Fig. 6A**, **Table S7**), but far fewer genes show altered expression in response to PQ in *unc-17* mutants compared with wild type (2,085 genes in wild type vs. 995 genes in *unc-17* mutant) (**Figs. 4B** and **6B,D**). Moreover, approximately 70% of the genes we identified to be upregulated in the wild-type transcriptional response to extended PQ exposure fail to upregulate in *unc-17* mutants (**Table S8**). Gene enrichment analysis showed that the stress response, transcription factor, and neuronal function categories are enriched amongst PQ-upregulated genes in *unc-17* mutants (**Fig. 6C**, **Table S7**), as in wild type. However, we noted substantial decreases in the number of upregulated genes contributing to each of these categories in *unc-17* mutants compared to the wild type. For example, the number of transcription factor genes significantly upregulated in the wild type are decreased by approximately 70% in *unc-17* mutants (126 in wild type compared with 35 in *unc-17* mutants). Genes in the neuronal function, metabolism, and extracellular material gene classes enriched in the wild-type transcriptional response to PQ are also reduced by approximately 70% in *unc-17* mutants. The stress response, cilia, and proteolysis/proteasome categories are exceptions to this pattern. Surprisingly, the stress response gene class is not as severely affected by *unc-17* mutation, ∼55% of the stress response genes we identified as upregulated in the wild-type transcriptional response to PQ are also upregulated in *unc-17* mutants. Conversely, the proteolysis/proteasome and cilia gene classes are more severely affected by disruption of ACh transmission. Only ∼20% of the proteolysis/proteasome genes and 13% of the cilia genes we found upregulated in the wild-type transcriptional response to PQ are also upregulated in *unc-17* mutants.

**Figure 6.**
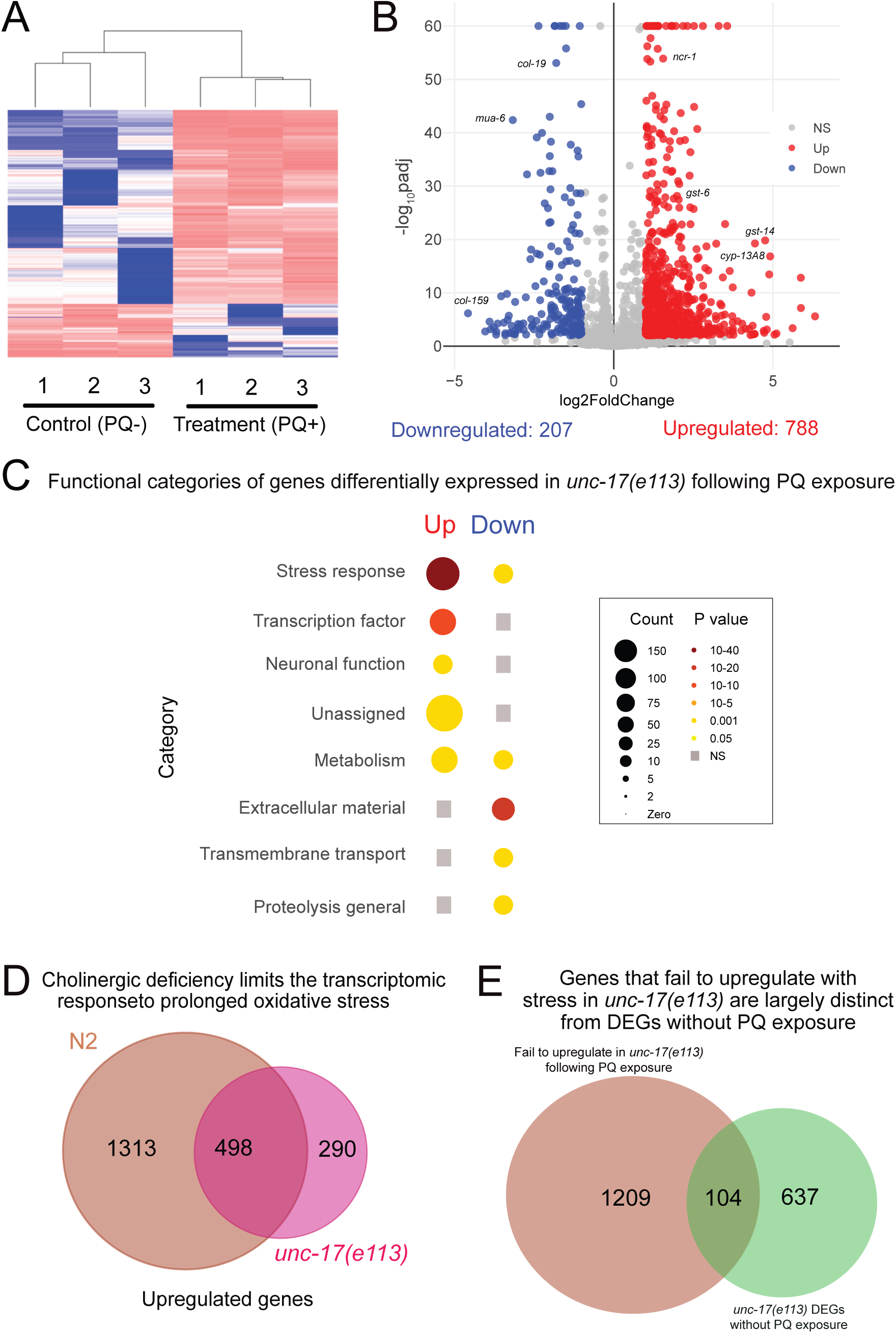
Acetylcholine deficiency blunts transcriptomic responses to oxidative stress Whole worm RNA sequencing data shows 48h continuous exposure to PQ (4 mM) does not change the transcriptomic landscape in *unc-17(e113)* mutants as severely as in wild type. (A) Heatmap shows clustering of treatment (PQ+) and control (PQ-) replicates for *unc-17(e113)*. Number of replicates: n=3 for both conditions. (B) Volcano plot (log_2_FoldChange, -log_10_padj) of genes differentially expressed in *unc-17(e113)* mutants in response to 48 hrs of PQ (4 mM) treatment compared to age-matched controls. Red: upregulated, Blue: Downregulated, Grey: not significantly different. Statistical cutoff for differential expression FoldChange>2, p_adj_<0.01, and False Discovery Rate (FDR) <0.01. (C) Functional categorization of differentially expressed genes (upregulated and downregulated) in *unc-17(e113)* mutants following 48 hr of PQ exposure. Grey squares indicate no significant enrichment in that category. Gene count and P-value scale as indicated. A larger radius corresponds to a higher number of genes in that category. A darker color corresponds to a more significant P value. (D) Venn diagram shows the intersection of upregulated genes in response to PQ for wild type (N2) (total 1811) and *unc-17(e113)* (total 788). 1,313 genes that are a part of the wild type transcriptional response to PQ fail to be upregulated in ACh-deficient *e113* mutants. (E) Venn diagram shows the intersection of PQ-responsive and ACh-dependent wild type genes compared with genes differentially expressed in *unc-17(e113)* versus wild type under control conditions without PQ treatment. 104 genes (14%) are shared between the two groups. Wild-type PQ-responsive genes that fail to upregulate in *unc-17* mutants following PQ treatment are largely distinct from those differentially expressed in *unc-17* without PQ treatment.

Interestingly, we also found that 290 additional genes, not included in the wild-type transcriptional response to PQ, are upregulated by PQ treatment in *unc-17* mutants (**Fig. 6D**). These additional genes largely fall into the same functional categories defined by the wild-type transcriptional response, with stress response and metabolism gene categories showing significant enrichment (**Tables S7 and S8**). These findings suggest the interesting possibility that activation of metabolic and stress pathways in *unc-17* mutants is achieved through transcriptional activation of alternate genes and relies on mechanisms less strongly dependent on ACh transmission. Notably, we found upregulation of very few additional genes in the proteolysis/proteasome class, perhaps suggesting that regulation of proteostasis through this gene class is more strongly dependent on ACh neurotransmission. Overall, we conclude that the transcriptional response to extended oxidative stress is strikingly blunted in *unc-17* mutants. Transcriptional activation of stress response genes is less severely compromised compared to other pathways comprising the wild-type transcriptional response, while transcriptional activation of proteolysis genes is most strongly impacted by disruption of ACh transmission.

To explore the requirements for cholinergic signaling in activation of an oxidative stress response versus other potential requirements that are independent of stress regulation, we also compared the transcriptional profiles of D3 adult wild type and *unc-17* mutants raised under control conditions (in the absence of PQ treatment). We identified 741 genes to be differentially expressed (429 upregulated, 312 downregulated) in *unc-17* mutants versus wild type (**Table S6**). Gene enrichment analysis showed that the stress response, neuronal function, muscle function, and transmembrane transport categories are significantly enriched amongst the differentially expressed gene set (**Table S6**). However, only ∼14% of the genes differentially expressed in *unc-17* mutants under control conditions were also upregulated as part of the *unc-17*-dependent transcriptional response to PQ (**Fig. 6E**, **Table S9**), suggesting there is little overlap between the transcriptional requirement for cholinergic signaling under control versus stress conditions. We conclude that cholinergic signaling is important for the mobilization of a transcriptional response to extended oxidative stress that is largely independent of potential roles for ACh signaling in organismal development.

### Deletion of the muscarinic acetylcholine receptor *gar-3* gene decreases survival during chronic oxidative stress

We next sought to investigate potential mechanisms by which ACh mediates transcriptional activation of an oxidant response program. Interestingly, we noted several genes involved in acetylcholine signaling were significantly upregulated in the wild-type transcriptional response to PQ. These included the M1/M3/M5 family G_q_ protein-coupled muscarinic acetylcholine receptor (mAChR) *gar-3* gene, the ionotropic nicotinic acetylcholine receptor (nAChR) subunit genes *lev-8* and *acr-12* (Petrash et al., 2013), and the *ric-3* gene that encodes a chaperone broadly required for the assembly and synaptic delivery of nicotinic acetylcholine receptors (Biala et al., 2009; Halevi et al., 2002) (**Fig. 7A**). Motivated by the transcriptomic studies, we asked whether impaired muscarinic or nicotinic signaling affected organismal vulnerability to chronic oxidative stress. Specifically, we quantified the survival of strains carrying deletion mutations in either *gar-3* or *ric-3* during chronic oxidative stress. While mutation of *ric-3* did not significantly alter survival during chronic PQ exposure, we observed a striking decrease in the survival of *gar-3* mutants in the presence of PQ (**Fig. 7B**). Further, the survival of *gar-3;ric-3* double mutants during PQ exposure was not appreciably different from that of *gar-3* single mutants (**Fig. 7C**). Notably, we did not find a significant decrease in the longevity of *gar-3* mutants under control conditions (PQ-), suggesting a potential protective role for cholinergic GAR-3 mAChR signaling against oxidative damage (**Fig. 7D**). Consistent with this, we found that the survival of *gar-3* mutants in the presence of PQ was decreased to a similar extent as observed for *unc-17* mutants (**Fig. 7E**). Moreover, the survival of *unc-17;gar-3* double mutants in the presence of PQ was not significantly decreased compared to either single mutant (**Fig. 7F**). Additionally, mutation of *gar-3* in combination with mutation of *gar-2*, another mAChR broadly expressed in the nervous system, did not enhance the survival deficits of *gar-3* single mutants (**Fig. 7F**). These results suggest that GAR-3 signaling may have a specialized role in promoting organismal resilience to oxidative stress; however, analysis of additional GPCR mutants (for instance *gar-1* mAChR and metabotropic glutamate receptors) is needed to draw this conclusion more firmly. Finally, transgenic expression of a wild-type GAR-3::YFP transgene that includes approximately 7.5 kb of the endogenous *gar-3* promoter region restored the survival of *gar-3* mutants on PQ to wild-type levels (**Fig. 7G**). Taken together, our findings demonstrate that the decreased survival of cholinergic-deficient mutants during chronic oxidant exposure occurs primarily due to deficits in cholinergic activation of the GAR-3 mAChR.

**Figure 7.**
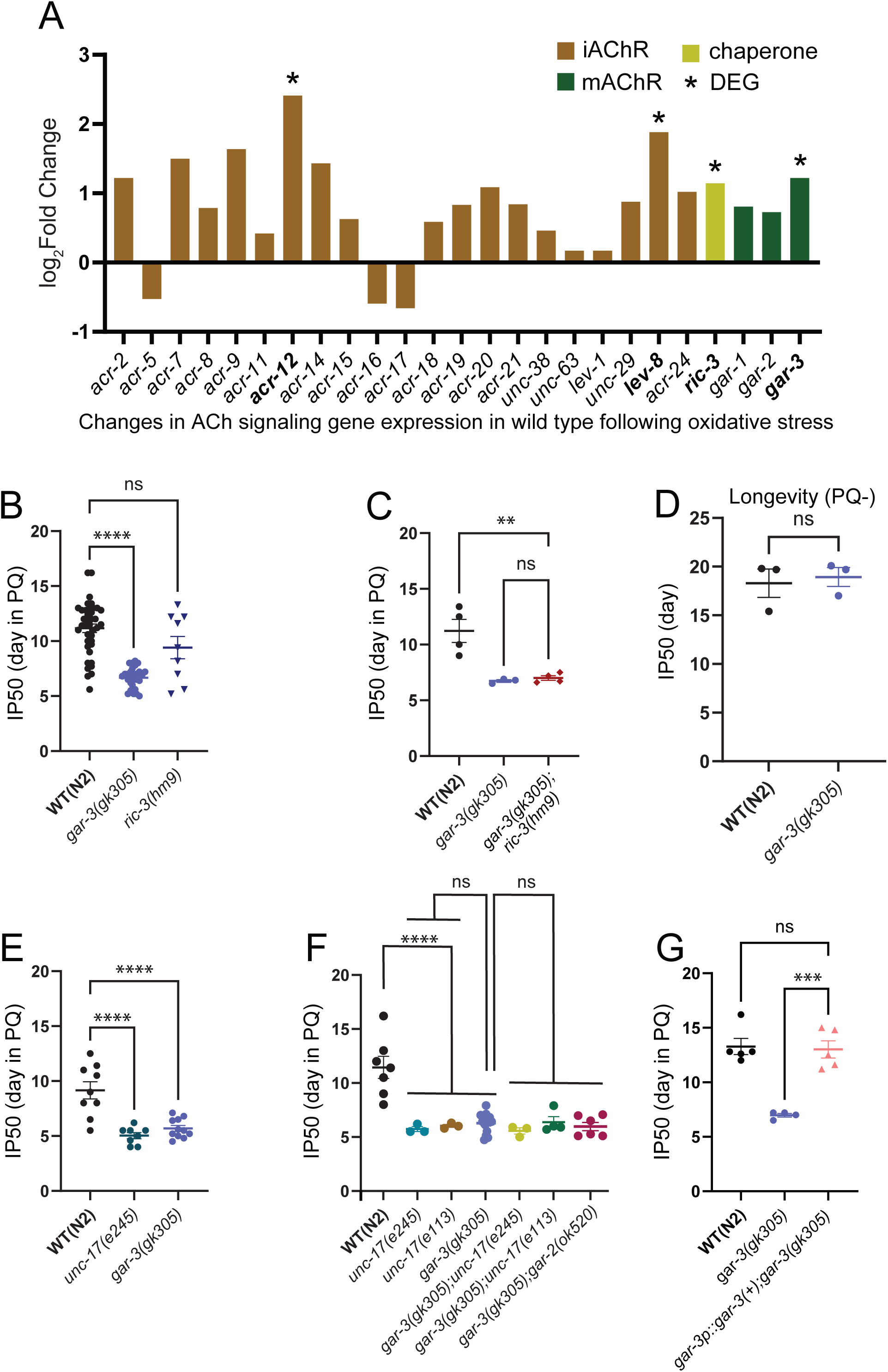
Cholinergic activation of GPCR signaling via mAChR GAR-3 promotes survival in the presence of oxidative stress (A) Bar graph showing ACh signaling genes upregulated in wild type in response to PQ. iAChR subunit *acr-12* and *lev-8*, chaperone *ric-3* and mAChR *gar-3* transcripts are significantly upregulated. The bar graph shows Log2Fold change value for ACh signaling gene transcripts (y axis). (B) Bar graph comparing IP50 measurements for wild type, *gar-3(gk305)* mAChR mutants and *ric-3(hm9)* chaperone mutants in the presence of PQ. Mutation of *gar-3* significantly reduces IP50 values compared to wild type, whereas the effect of *ric-3(hm9)* is variable. Each data point represents an independent trial. Bars indicate mean ±SEM. *\*\*\*\*p<0.0001, gar-3(gk305); p= 0.0531(ns), ric-3(hm9).* One-way ANOVA with Dunnett’s multiple comparisons test. (C) Bar graph comparing IP50 measurements for wild type, *gar-3(gk305)* mutants and *gar-3*;*ric-3* double mutants in the presence of PQ. The IP50 of *gar-3*;*ric-3* double mutants is similar to that of *gar-3(gk305)* single mutants. Each data point represents one independent trial. Bars indicate mean ± SEM. *\*\*p= 0.0044,* WT (N2) *vs.gar-3(gk305);ric-3(hm9); p=0.0961(ns) for gar-3(gk305) vs. gar-3(gk305);ric-3(hm9).* One-way ANOVA with Tukey’s multiple comparisons test. Number of trials: n=4, N2; n=3, *gar-3(gk305)*; n=4, *gar-3(gk305);ric-3(hm9)*. (D) Bar graph comparing IP50 measurements for wild type and *gar-3(gk305)* mutants under control condition in the absence of PQ. *gar-3* mutants do not show an appreciable reduction in longevity compared to wild type under control (PQ-) conditions. Each data point represents one independent trial. Bars indicate mean ± SEM. *ns,* unpaired two-tailed t-test with Welch’s correction. Number of trials: n= 3. (E) Bar graph comparing IP50 measurements for wild type, *unc-17(e245)* and *gar-3(gk305)* mutants in the presence of PQ. Mutation of *unc-17* or *gar-3* reduces survival to a similar extent in the presence of PQ. Each data point represents one independent trial. Bars indicate mean ± SEM. *\*\*\*\*p<0.0001*, one-way ANOVA with Dunnett’s multiple comparisons test. Number of trials: n= 9, N2; 8, *unc-17(e245)*; 11, *gar-3(gk305)*. (F) Bar graph comparing IP50 measurements for wild type, *unc-17(e245)*, *unc-17(e113)*, *gar-3(gk305)* and double mutant combinations in the presence of PQ. Combined mutation of *gar-3* and *unc-17* elicits a similar reduction in IP50 to that of either single mutant, providing evidence that *unc-17* and *gar-3* may act in the same pathway. In addition, combined mutation of *gar-3* and *gar-2* elicits a similar reduction in IP50 to that of *gar-3* single mutants. Each data point represents one independent trial. Bars indicate mean ± SEM. \*\*\*\**p=0.0001*, WT vs. *unc-17(e245)*; WT vs. *unc-17(e113)*; WT vs. *gar-3(gk305).* One-way ANOVA with Tukey’s multiple comparisons test. Number of trials: n≥3. (G) Bar graph comparing IP50 measurements for wild type, *gar-3(gk305)* mutants and transgenic *gar-3* mutants expressing *gar-3(+)*. Expression of wild type *gar-3* using the native *gar-3* promoter region normalizes the survival of *gar-3* mutants in the presence of PQ. Each data point is one independent trial. Bars indicate mean ± SEM. ***p=0.0002, *gar-3(gk305)* vs. *gar-3p::gar-3(+*);*gk305.* One-way ANOVA with Tukey’s multiple comparisons test. Number of trials: n≥4.

To better define potential mechanisms for GAR-3 regulation of antioxidant responses, we next sought to analyze the cellular expression of the *gar-3* gene. First, analyzed *gar-3* expression using a genome-edited reporter strain, *gar-3(syb9584)*, in which the SL2 splice leader and GFP coding sequence are inserted into the *gar-3* genomic locus. We noted *gar-3* expression in the nerve ring and motor neurons of the ventral nerve cord, as well as in body wall muscles and pharynx (**Fig. 8A-C**). Using strains that co-express *gar-3::SL2::GFP* with red fluorescent (mCherry) reporters, we determined that *gar-3* is expressed in both cholinergic (*acr-2p::mCherry*) and GABAergic (*unc-47p::mCherry*) ventral cord motor neurons (**Fig. 8D,E**), consistent with prior single neuron RNA-seq findings (Smith et al., 2024; Taylor et al., 2021). In addition, we analyzed the expression of a *gar-3p::GFP* transcriptional reporter containing the same promoter region as used for rescue. We found that this reporter broadly recapitulated the expression pattern of the *gar-3::SL2::GFP* strain, including expression in both muscles and ventral cord motor neurons, particularly cholinergic ventral cord motor neurons as previously reported (**Fig. S3A-C**) (Chan et al., 2013). To assess the cellular requirements for rescue of the reduced survival phenotype, we expressed wild type *gar-3* cDNA in either cholinergic motor neurons or muscles of *gar-3* mutants. We found that expression of *gar-3* in either cell type provided weak, partial rescue of the *gar-3* mutant survival phenotype as assessed from survival curves; however, this rescue was not apparent from the more stringent IP50 measurement (**Figs. 8F,G** **and S3D**). We cautiously interpret these results to suggest that GAR-3 signaling in both tissues may contribute toward stress resilience, though we can not rule out contributions from other cell types.

**Figure 8.**
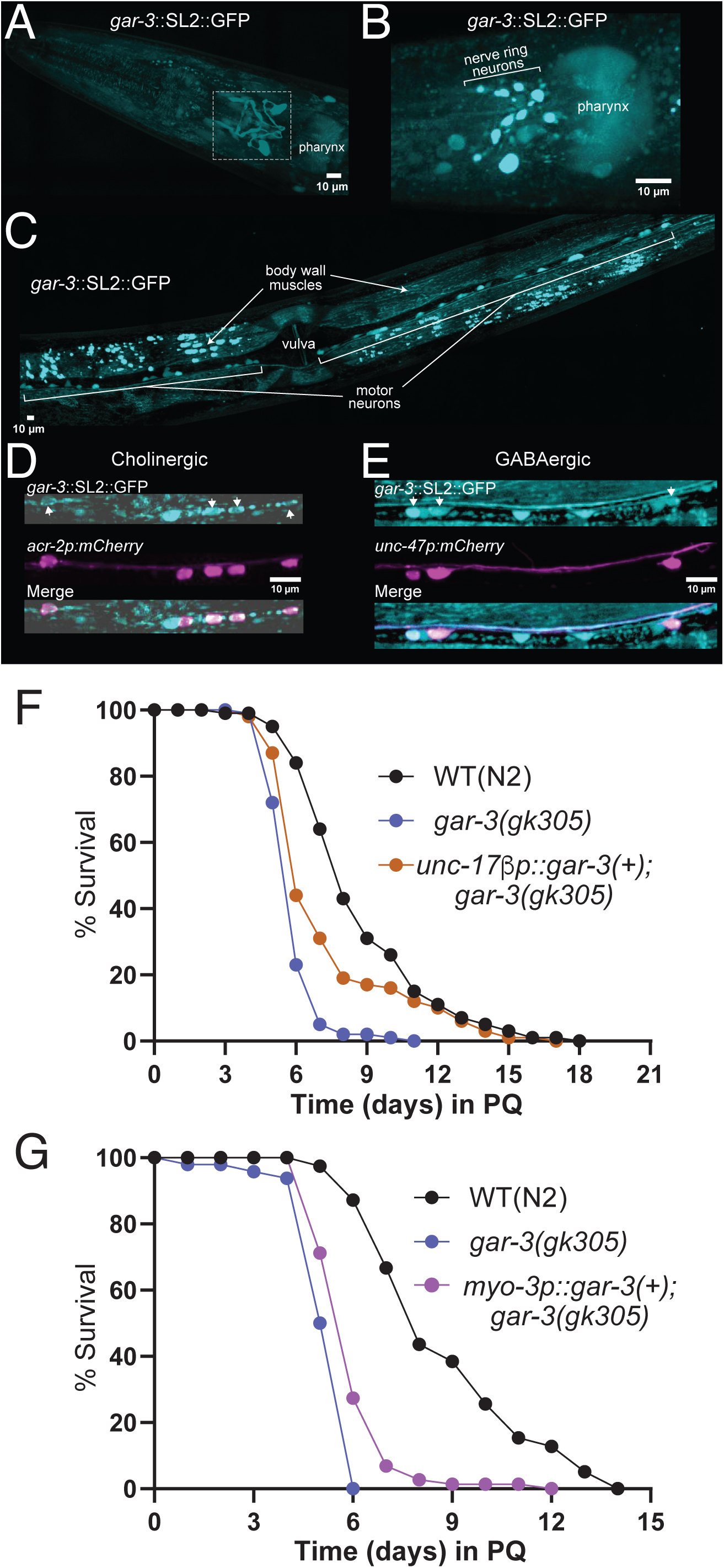
*gar-3* expression and tissue-specific rescue **(A)** Confocal maximum intensity projection of the head region of an adult animal expressing genome-inserted *gar-3::SL2::GFP*. Note expression in nerve ring processes (white box) and in the pharynx. Anterior is to the left in all panels. Scale, 10 µm. **(B)** Confocal maximum intensity projection showing zoomed-in view of *gar-3* expression in nerve ring neuronal cell bodies. Scale, 10 µm. **(C)** Confocal maximum intensity projection of the midbody region showing expression in ventral nerve cord motor neurons and body wall muscles. Scale, 10 µm. **(D)** Confocal maximum intensity projections of a segment of the ventral nerve cord showing expression of *gar-3*::SL2::GFP in cholinergic motor neurons labeled by *acr-2p::mCherry*. White arrows indicate *gar-3* expressing cholinergic neurons. Scale bar, 10 μm. **(E)** Confocal maximum intensity projections of a segment of the ventral nerve cord showing expression of *gar-3*::SL2::GFP in GABAergic motor neurons labeled by *unc-47p::mCherry*. White arrows indicate *gar-3* expressing GABAergic neurons. Scale bar, 10 μm. **(F)** Kaplan-Meier survival curves comparing wild type, *gar-3* mutant and ACh motor neuron rescue strains. Specific expression of wild type *gar-3* cDNA (10 ng/µl) in ACh motor neurons of *gar-3* mutants partially rescues the decreased survival of *gar-3* mutants in PQ. Kaplan-Meier curves are cumulative of ≥4 trials. *\*\*p= 1.5E-03*, *gar-3(gk305)* vs. *unc-17βp::gar-3(+*);*gar-3(gk305),* Fisher’s exact test at 50% of survival. Total number of animals: n=105, N2; 97, *gar-3(gk305)*; 96, *unc-17βp::gar-3(+*);*gar-3(gk305)*. **(G)** Kaplan-Meier survival curves comparing wild type, *gar-3* mutant and muscle-specific rescue strains. Specific expression of wild type *gar-3* cDNA muscles of *gar-3* mutants weakly rescues the decreased survival of *gar-3* mutants in PQ. Kaplan-Meier curves are cumulative of X trials. *\*\*\*\*p= 5.2e-06*, *gar-3(gk305)* vs. *myo-3p::gar-3(+*);*gar-3(gk305)*, Fisher’s exact test at 50% of survival. Total number of animals: n=45, N2; 48, *gar-3(gk305)*; 75, *myo-3p::gar-3(+*);*gk305*.

### Deletion of *gar-3* severely weakens the transcriptional response to extended oxidative stress

To determine the requirement for GAR-3 mAChR signaling in promoting the ACh-dependent transcriptional response to chronic oxidant exposure, we performed bulk RNA sequencing of day-3 adult *gar-3* deletion mutants under control conditions or following 48 hr of 4 mM PQ exposure. We observed a significant transcriptional response to extended oxidative stress in *gar-3* mutants (**Fig. 9A**), but, as for *unc-17* mutants, noted far fewer PQ stress-responsive genes in *gar-3* mutants compared with wild type (2,085 genes in wild type vs. 1,004 genes in *gar-3* mutants) (**Figs. 4B**, **9B**). We identified 868 upregulated and 136 downregulated genes in *gar-3* mutants (**Fig. 9B**). Similar to our findings for *unc-17* mutants above, approximately 70% of the genes we identified to be upregulated in the wild-type transcriptional response to extended PQ exposure failed to upregulate in *gar-3* mutants (**Tables S1 and S10**). To provide additional confirmation of these findings, we pursued parallel strategies to investigate the expression of two candidates identified by RNA-seq as upregulated in response to prolonged oxidative stress in the wild type, but not in *unc-17* or *gar-3* mutants. Consistent with our RNA-seq findings, we found that a *nhr-185pr::GFP* transcriptional reporter was upregulated in the wild type by PQ treatment, mainly in the pharynx and anterior intestine (**Fig. S4A**). By quantitative RT-PCR, we found that expression of the *fbxa-73* gene was upregulated in the wild type by PQ treatment, providing independent evidence supporting our RNA-seq findings (**Fig. S4B**).

**Figure 9.**
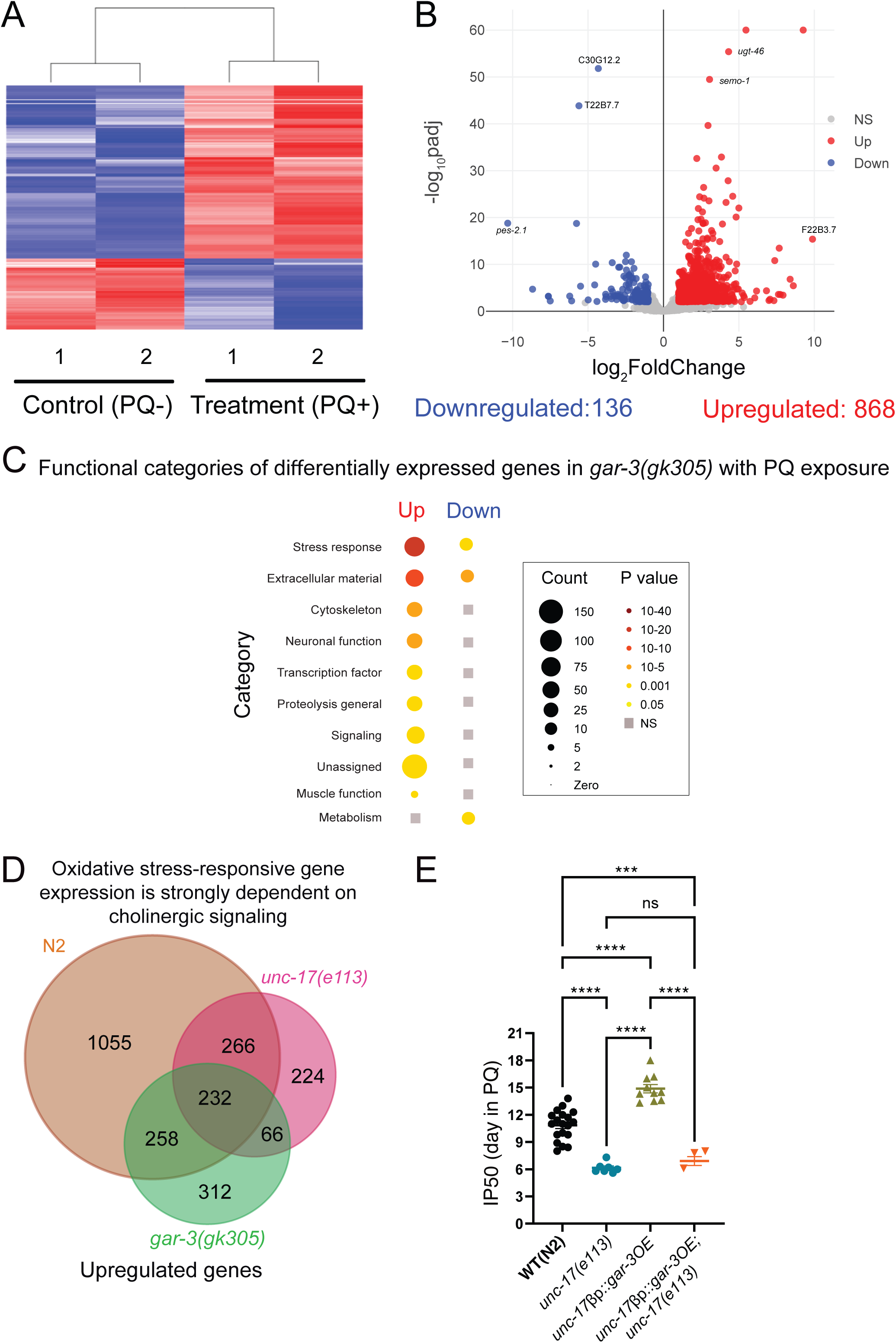
Deficiency of cholinergic signaling via GAR-3 blunts transcriptomic responses to oxidative stress (A) Heatmap shows clustering of treatment (PQ+) and control (PQ-) replicates for *gar-3(gk305)*. n=2 for both conditions. (B) Volcano plot (log_2_FoldChange, -log_10_padj) of genes differentially expressed in *gar-3(gk305)* mutants in response to 48 hr of PQ (4 mM) treatment compared to age-matched controls. Red: upregulated, Blue: Downregulated, Grey: not significantly different. Statistical cutoff for differential expression FoldChange>2, P_adj_<0.01 and False Discovery Rate (FDR) <0.01. (C) Functional categorization of differentially expressed genes (upregulated and downregulated) in *gar-3(gk305)* mutants in the presence of extended PQ exposure. Grey squares indicate no significant enrichment in that category. Gene count and P-value scales as indicated. A larger radius corresponds to a higher number of genes in that category. A darker color corresponds to a more significant P value. (D) Venn diagram showing the intersection of upregulated genes in wild type (total 1811), *unc-17(e113)* (total 788), and *gar-3(gk305)* (total 868) in response to PQ. 1055 genes that are upregulated in the wild transcriptional response to PQ fail to upregulate in both ACh-deficient *unc-17* and *gar-3* mAChR mutants. (E) Bar graph comparing IP50 measurements for wild type, *unc-17(e113)*, transgenic wild type animals overexpressing *gar-3* in cholinergic motor neurons (*unc-17βp::gar-3* OE) and transgenic *unc-17* mutants overexpressing *gar-3* in cholinergic motor neurons (*unc-17βp::gar-3(+);unc-17* ) in the presence of PQ. Cholinergic overexpression of *gar-3* significantly increases survival compared to wild type. Mutation of *unc-17* reverts survival to the level of *unc-17(e113)* mutants. Each data point is one independent trial. Bars indicate mean ±SEM. *\*\*\*\*p<0.0001,* WT(N2) *vs. unc-17(e113),* WT(N2) *vs. unc-17βp::gar-3OE, unc-17(e113) vs. unc-17βp::gar-3OE,* and *unc-17βp::gar-3OE vs. unc-17βp::gar-3OE;unc-17. ***p=0.0005,* WT(N2) *vs. unc-17βp::gar-3(+);unc-17.* One-way ANOVA with Tukey’s multiple comparisons test. Number of trials: n=19, WT(N2); 7, *unc-17(e113); 10, unc-17βp::gar-3OE; 3, unc-17βp::gar-3(+);unc-17*.

Notably, the stress response, transcription factor, and neuronal function categories were significantly enriched amongst PQ-upregulated genes in *gar-3* mutants (**Fig. 9C**, **Table S10**).

Similar to *unc-17* mutants, the proteolysis proteasome and cilia categories were not represented amongst the enriched gene categories (**Figs. 6C, 9C**). Roughly 250 genes upregulated in the wild-type transcriptional response to PQ were also upregulated in both *unc-17(e113)* and *gar-3(gk305)* mutant transcriptional responses (**Fig. 9D, Table S12**). These genes include oxidative stress response genes such as *gst-4*, *gst-14,* and the heavy metal stress response gene *cdr-1*, known to be regulated by SKN-1, suggesting SKN-1-dependent upregulation of detoxification genes occurs largely independent of cholinergic signaling through GAR-3.

We also identified 1,055 genes that were upregulated by PQ exposure in wild type, but failed to upregulate in both *unc-17(e113)* and *gar-3(gk305)* mutants (**Table S11**). We propose that transcriptional activation of these genes during extended oxidative stress requires cholinergic neurotransmission via GAR-3. Therefore, we expect enriched categories amongst this gene set will identify stress-responsive pathways that are regulated through muscarinic activation. We identified the antioxidant gene *ctl-1*, multiple detoxification genes (*gst-20*, *ugt-34*, *cyp-13A6*), and C-type lectins (*clec-83*, *clec-160*) within the group of stress-responsive genes that required cholinergic signaling and GAR-3 activation. In addition, the *nhr-49*-dependent, stress-responsive *fmo-2* gene (Doering et al., 2022; Huang et al., 2021; Leiser et al., 2015; Wani et al., 2021) failed to upregulate in both *unc-17* and *gar-3* mutants, suggesting transcriptional responses to oxidative stress mediated through *nhr-49* are at least partially dependent on ACh signaling and GAR-3 activation.

As the proteolysis proteosome category of genes appeared strongly dependent on ACh signaling and GAR-3 activation, we also further investigated genes in this group. Proteolysis proteasome genes that are upregulated by PQ exclusively in wild type primarily consist of E3 domain F box genes (e.g. *fbxa-73*, *fbxb-22*, *fbxc-34*). F-box proteins act as substrate adaptors for the S-phase kinase-associated protein 1 (SKP1)-cullin 1 (CUL1)-F-box protein (SCF) ubiquitin ligase complexes, which mediate proteasomal degradation of a diverse range of regulatory proteins. Our results therefore suggest that cholinergic signaling through the GAR-3 mAChR may regulate proteostasis during oxidative stress through upregulation of ubiquitin-dependent protein degradation.

### *gar-3* overexpression in cholinergic motor neurons increases survival during chronic oxidative stress

As our expression studies indicated significant *gar-3* expression in ventral cord motor neurons and we obtained partial rescue with motor neuron-specific *gar-3* expression, we next asked whether overexpression of *gar-3* may be sufficient to offer protection from oxidative damage and increase organismal survival during chronic PQ exposure. While pan-neuronal *gar-3* overexpression proved lethal, we were able to obtain viable strains overexpressing *gar-3* in a subset of cholinergic neurons, including ventral cord motor neurons. Transgenic animals injected with the *unc-17βp::gar-3* transgene (25 ng/µL) displayed strikingly increased survival in the presence of 4 mM PQ compared to either *gar-3* mutants or wild type (**Fig. 9E**). In contrast, overexpression of *gar-3* in cholinergic motor neurons of *unc-17* mutants did not extend survival. Indeed, transgenic *unc-17* mutants expressing *unc-17βp::gar-3* survived similarly to *unc-17* single mutants during chronic PQ exposure. Thus *gar-3* overexpression in cholinergic motor neurons is sufficient to elicit protection from oxidative stress as indicated by increased survival to PQ, and these protective effects are dependent on vesicular release of ACh.

## DISCUSSION

Our studies reveal new mechanistic insights into neural regulation of organismal stress responses. Using temporally defined periods of neuronal silencing in combination with analyses of genetic mutants deficient for specific neurotransmitters, we demonstrate that neuronal activity prior to and in the early stages of stress exposure are each important for the activation of stress-responsive transcriptional programs that promote organismal resilience in the face of extended oxidative stress. Cholinergic transmission is key for mobilizing this transcriptional stress response program. Our transcriptomic and survival analyses provide evidence that cholinergic GAR-3 activation is particularly important for stress-responsive upregulation of genes important for ubiquitin-dependent protein degradation, offering insight into a potential circuit for inter-tissue control of proteostasis. Finally, we show that overexpression of GAR-3 in cholinergic motor neurons confers enhanced survival to chronic oxidative stress, perhaps pointing to cholinergic motor neurons as a critical cellular site of action for the regulation of antioxidant responses.

Prior work has suggested that activation of extrasynaptic GAR-3 mAChRs by humoral ACh promotes presynaptic release from cholinergic motor neurons (Chan et al., 2012; Chan et al., 2013). Based on this finding and our data, we hypothesize that elevated synaptic release from cholinergic motor neurons through GAR-3 activation may be important for coordinating an organismal response to chronic oxidative stress. Prior work implicates muscarinic signaling in the activation of host defenses against pathogen infection, perhaps suggesting interesting mechanistic parallels between these two protective responses (Goswamy and Irazoqui, 2021; Labed et al., 2018). Notably, cholinergic neurons are among those reported to be most vulnerable to degeneration during the early stages of Alzheimer’s disease (AD) (Chen et al., 2022; Davies and Maloney, 1976; Mufson et al., 2008). Our findings may point toward interesting links between cholinergic signaling, muscarinic activation, and oxidative stress-mediated degeneration during AD.

### Basal neural activity may promote organismal survival through neurohormesis

Our pan-neuronal silencing experiments provided evidence for neuronal activity-dependent regulation of a protective response to oxidative stress. Our follow-up genetic studies implicated both glutamatergic and cholinergic systems in protective responses to oxidative stress. The effects of pan-neuronal silencing on survival during long-term PQ exposure may therefore be derived solely from cholinergic neurons, glutamatergic neurons, or a combination of both neuronal populations. Additional neuron-specific silencing experiments will be required to distinguish between these possibilities. Interestingly, we also found that neuronal silencing either prior to or during the early stages of oxidative stress each decrease organismal survival, suggesting that neural activity may have roles both in acute responses to oxidative stress and in equipping the organism to mount an effective protective response. Neural activity and ROS production are intimately associated due to the high energetic cost of maintaining neuronal excitability (Harris et al., 2012; Hongpaisan et al., 2004). Prior work has demonstrated that neuronal adaptation to harsh environments is elicited through prior exposure to sub-lethal doses of particular stressors. In particular, neurons appear to retain a ‘memory’ from prior exposure to a low dosage of a stressor that enables the animals to defend against that stressor in high doses, a phenomenon known as ‘neurohormesis’ (Harris et al., 2012; Hongpaisan et al., 2004; Mattson and Cheng, 2006; Robertson, 2004). An example of such ‘preconditioning’ is found in the protection of rat cerebellar granule cells from glutamate-mediated excitotoxicity where low level stimulation of NMDA receptors elicits activation of nuclear-factor kappaB (NF-kB), a transcription factor involved in protective responses to stress (Jiang et al., 2003). Notably, neurohormesis responses are often mediated through transcriptional regulation (Jeong et al., 2006; Liu et al., 2005; West et al., 2001). Given the broad transcriptional response to extended PQ exposure we observed, our finding that silencing prior to PQ treatment decreases survival may suggest a similar form of neuronal priming. We speculate that neural activity may induce a modest increase in ROS which primes a protective transcriptional response, preparing the animals to defend against future oxidative stress.

### Acute and chronic oxidative stress elicit distinct molecular responses

Prior work indicated that a deficiency in acetylcholine neurotransmission either had no effect (Jia and Sieburth, 2021) or was protective (Cornell et al., 2024; Jia and Sieburth, 2021) in cases of acute oxidant exposure. Surprisingly, we found that ACh-deficient mutants had severely decreased survival to chronic oxidative stress, perhaps pointing to interesting mechanistic differences in the transcriptional programs initiated by acute versus chronic stress and how they are regulated by the nervous system. Though comparisons across these studies are complicated by the fact that different oxidants are known to elicit distinct transcriptional responses (Goh et al., 2018; Oliveira et al., 2009; Wu et al., 2016), we noted interesting differences between our transcriptome analyses of extended stress versus previous analyses of acute stress responses. While many of the same core antioxidant genes and detoxification gene classes are upregulated in both cases, we observed a much broader transcriptional response to extended exposure that encompassed additional gene categories. We suggest that cholinergic signaling recruits additional stress-responsive genes that are important for coping with the impacts of extended stress. The differences between transcriptional responses to acute and chronic oxidant exposure may have important implications for understanding organismal responses to different classes of stressors, for example, acute exposure to an environmental toxin versus extended stress associated with chronic disease.

### Cholinergic mAChR activation is required for upregulation of genes important for ubiquitin-dependent protein degradation

Our gene enrichment analysis demonstrated that several gene categories, including proteolysis/proteasome, transmembrane transport, and cilia gene classes, which were enriched in the wild-type transcriptional response were no longer enriched in the *unc-17* or *gar-3* transcriptional responses. The loss of enrichment for the proteolysis/proteasome gene class, particularly E3 ubiquitin ligases, is remarkable because this gene category was one of the three most strongly upregulated in the wild-type gene set. This suggests a more stringent requirement for cholinergic transmission in the upregulation of proteolysis/proteasome genes in response to extended stress. Maintaining protein homeostasis is critical for organismal health (Koga et al., 2011). Like other eukaryotes, *C. elegans* depend on the ubiquitin-proteasome system for selective and efficient degradation or recycling of damaged or unneeded proteins (Papaevgeniou and Chondrogianni, 2014; Tanaka, 2009). The ACh-dependent upregulation of E3 ubiquitin ligases observed in our transcriptomic and quantitative RT-PCR (*fbxa-73*) studies raises an intriguing link between cholinergic transmission and the regulation of proteostasis through the ubiquitin-proteasome system. Recent evidence has suggested interesting links between cholinergic neurotransmission and proteostasis in other contexts. For instance, recent work showed that depletion of *C. elegans* BAZ-2, ortholog of *BAZ2B* (bromodomain adjacent to zinc finger domain 2B), reduced toxicity and aggregation of aggregation-prone peptides associated with AD and poly-Q diseases (Gallrein et al., 2023). BAZ-2 is a negative regulator of ACh metabolism and signaling, suggesting ACh signaling promotes proteostasis. In support of this conclusion, the authors showed that acetylcholine treatment can augment proteostasis by induction of both the endoplasmic reticulum unfolded protein response and the ubiquitin-proteasome system. Similarly, increased cholinergic signaling onto *C. elegans* muscles induces calcium-dependent activation of the stress-responsive transcription factor HSF-1 and increased expression of cytoplasmic chaperones, resulting in suppression of protein misfolding (Silva et al., 2013). Our transcriptomic and survival data suggest that cholinergic signaling also regulates proteostasis during prolonged oxidative stress.

### Cholinergic activation of the GAR-3 mAChR promotes organismal survival

Our transcriptome analysis, survival assays and rescue experiments identified a critical role for GAR-3 in the mobilization of the transcriptional response to extended oxidative stress. Our analysis of the *nhr-185* transcriptional reporter showed stress-responsive transcriptional upregulation of *nhr-185* occurs largely in the pharynx and anterior intestine. *gar-3* overexpression in cholinergic motor neurons increased survival during oxidative stress in an ACh-dependent manner. Collectively, these results provide evidence for neuronal muscarinic activation of a protective antioxidant transcriptional response that is mobilized, at least in part, in peripheral tissues. GAR-3 is most similar to vertebrate M1/M3/M5 mAChR (Eglen, 2006; Kato et al., 2021). Previous studies of a variety of mammalian cell types have provided evidence that activation of the muscarinic M3 receptor can reduce oxidative stress (De Sarno et al., 2003; De Sarno et al., 2005; Frinchi et al., 2019; Giordano et al., 2009). Our work demonstrates that muscarinic activation can promote organism-wide antioxidant defenses and suggests that muscarinic control of antioxidant responses is evolutionarily conserved. It is interesting to speculate that similar muscarinic signaling pathways may coordinate antioxidant responses across cell types in the brain, for example between neurons and glia. Therapeutic approaches targeting muscarinic receptors may therefore offer a path to combat neurodegenerative disease and other pathological conditions resulting from extended oxidative stress.

## MATERIAL AND METHODS

### *C. elegans* strains and genetics

All *C. elegans* strains are derivatives of the N2 Bristol strain (wild type) and maintained under standard conditions at 20°C on nematode growth media plates (NGM) seeded with the *E. coli* strain OP50. A complete strain list is provided in **Table S13.** Transgenic strains were generated by microinjection of plasmids or PCR products into the gonad of young hermaphrodites (Mello et al., 1991).

### Molecular biology

Plasmids were constructed using the two-slot Gateway Cloning system (Invitrogen) and confirmed by restriction digestion and/or sequencing as appropriate. All plasmids and primers used in this work are described in Table 2 and Table 3 respectively.

pKB1 (*punc-17βp::gar-3 cDNA*) was created by recombining pENTR-*unc-17β* with pDEST-254 (*gar-3* cDNA vector construct). *gar-3* cDNA was amplified from wild-type RNA with Superscript 3 (Invitrogen) using primers OMF2777 (forward: 5’ATTAGGTACCATGCAGTCCTCTTCGTTGG3’) and OMF2778 (5’ GCTAGCCGGCCTAGTTGCGTCGGACATA3’) and ligated into Kpn1/MgoMIV digested pDEST-16 to generate pDEST-254. pKB1 was injected into wild type (20 ng/µl) for overexpression or *gar-3(gk305)* (10 ng/µl) for rescue. pLM18 (*myo-3p::gar-3* cDNA) was generated by recombination of pENTR-3’-myo-3 with pDest-254 and injected (10 ng/µl) into *gar-3(gk305)* for rescue. pKB5 (*gar-3p::GFPnovo2*) was generated by recombining pENTR-83 and pDEST-187. An 8.5 kb region immediately upstream of *gar-3* ATG (initiation codon) was amplified from genomic DNA using primers OMF3151 (5’CACCCGAGGGTGTTGCTCATTTCTAAACA3’) and OMF3152 (5’CACCCCTCTCGTCTGTGGTGATCCTGTAA3’) and ligated into pENTR-D-TOPO to create pENTR-83 (Chan et al., 2013). pDEST-187 was created by inserting a ccdB cassette into pSM.GFPnovo2 (gift from K. Mizomoto). The *gar-3p*:*gar-3::YFP::unc-54* plasmid (PYL8) was a gift from R. Garcia.

### Chronic PQ exposure assay

Chronic PQ exposure assays were performed as described previously (Senchuk et al., 2017). PQ (Sigma-Aldrich CAS number: 75365-73-0 and Thermo Scientific CAS number: 1910-42-5) and FUDR (Sigma-Aldrich, CAS number: 50-91-9) were dissolved in dH2O and added to NGM to final concentrations of 4 mM and 100 µM respectively. The experimental PQ concentration was selected based on prior work demonstrating its effectiveness as an oxidant in prolonged stress studies (Dues et al., 2016; Schaar et al., 2015; Senchuk et al., 2017; Van Raamsdonk and Hekimi, 2012). Briefly, animals were raised continuously at 20°C and grown on NGM plates until adult Day 1 stage, after which they were transferred onto plates containing 4 mM PQ and 100 µM FUDR. These plates were seeded with 10x concentrated *E. coli* OP50 culture and roughly 30 Day 1 adult animals were placed on each plate. The number of animals that died on each plate was recorded each day and surviving animals were transferred to new plates at regular intervals until no surviving animals remained. Animals were scored as dead if they did not respond to nose or tail touch. Unless indicated otherwise, all chronic PQ exposure assays were repeated for at least 3 individual trials with at least 20 animals per trial.

### Statistical analyses

Kaplan-Meier survival curves were generated using the OASIS opensource platform (https://sbi.postech.ac.kr/oasis/introduction/) (Yang et al., 2011). Fisher’s exact test was performed to determine the statistical significance of the differences in survival between experimental strains. Unless otherwise indicated, the p-values at 50% survival are reported for Fisher’s exact tests. For the Initial Population 50 (IP50) calculation, Kaplan-Meier survival curves were made for each trial with each genotype. The time taken to reach 50% of the initial population was calculated using the Kaplan-Meier estimator. For determining the statistical significance of the difference of IP50 values of different strains, the IP50 values were subjected to One-Way ANOVA with Dunnett’s or Tukey’s multiple comparisons test. Graphs were plotted using GraphPad Prism.

### Longevity assay

For longevity assays animals were staged in a similar way as the chronic PQ exposure assay described earlier. Instead of plates containing PQ, day 1 adult animals were placed on NGM agar plates containing FUDR (Sigma-Aldrich) to a final concentration of 100 µM. Plates were seeded with 10x concentrated OP50. Surviving animals were transferred to fresh plates at regular intervals. The number of surviving animals was recorded as described earlier. Kaplan-Meier survival curves and statistical tests were performed as described above.

### Protein carbonyl measurement

The Oxyblot^TM^ immunoblot approach was used to determine oxidative damage based on the detection of 2,4-dinitrophenylhydrazine derivatized carbonyls in protein(Goudeau and Aguilaniu, 2010; Nasrallah et al., 2023). For age synchronization, large populations (2000-3000) were grown on 10x concentrated OP50 until gravid adulthood and bleached to collect eggs. After repeated washes with M9, the eggs were plated on NGM with 10X concentrated OP50 and grown at 20°C. Around 40 hours post hatch (L4 stage), 25 µl of 100 mM FUDR was added to the plates. Day 1 adult animals were transferred to 4 mM PQ or control plates the following day for a period of 48 hours. Treated animals (PQ +/-) were washed free of bacteria with M9 and resuspended in 1x PBS. Worm pellets were snap frozen with liquid N_2_ and stored at -80°C. Samples were thawed in RIPA lysis buffer (ThermoFisher, #89900), and Halt protease inhibitor (ThermoFisher, #78429) and sonicated by Teflon homogenizer. The protein-containing supernatant was collected following centrifugation. Protein concentrations were determined by the DC protein assay (DC^TM^ protein assay kit II 5000112), using BSA as a standard. The Oxyblot^TM^ protein oxidation detection kit (Sigma-Aldrich S7150) was used to detect DNPH derivatized carbonyl protein per manufacturer instructions. Protein samples were incubated at room temperature with SDS and 2,4-dinitrophenylhydrazine (DNPH) for 15 mins. The samples were neutralized and 15-20 μg of the sample protein solution were loaded for electrophoresis using NuPage 4–12% gels (Life Technologies), transferred to nitrocellulose membranes (Life Technologies) under vacuum pressure, blocked with 5% fat-free milk in TBST and probed with a rabbit-anti DNPH polyclonal primary antibody and goat anti-rabbit IgG (HRP conjugated) antibody. Protein bands were visualized using a ChemiDoc MP Imaging System (Bio-Rad Laboratories, Inc.) and Supersignal^TM^ West Femto (Thermo Scientific 34096) was used as the chemiluminescent substrate. DNPH derivatized protein intensity was normalized against tubulin and quantified using Fiji imaging software (Schindelin et al., 2012). Statistical significance was determined using an unpaired two tailed t-test with Welch’s correction. Plots were made with GraphPad Prism.

### NAC treatment

*N*-acetylcysteine (NAC) (Sigma -Aldrich, CAS number: 616-91-1) was prepared in dH_2_O and added into NGM to a final concentration of 9 mM following a published protocol (Desjardins et al., 2017). Similarly, to prepare NAC+PQ plates, NAC and PQ were added to NGM to final concentrations of 9 mM and 4 mM, respectively. Plates were seeded with 10x *E. coli* OP50 and Day 1 adult animals were transferred to plates containing either NAC or NAC+PQ. Survival curves and statistical comparisons were performed as described above.

### Neuronal silencing and survival assay

Neuronal silencing was performed following the published protocol (Pokala et al., 2014). Briefly, transgenic animals expressing *kyEx4571* (*tag168p::HisCl1::SL2::GFP; myo-3p::mCherry)* were raised on plates containing 10 mM histamine dihydrochloride (Sigma-Aldrich, CAS number 56-92-8) for 24 hours at 20°C and exposed to 4 mM PQ at the indicated time period or maintained on control plates. Kaplan-Meier survival curves and IP50 calculations were performed as previously. Statistical comparisons were performed using two-tailed unpaired t-test with Welch’s correction.

### Whole-worm bulk RNA sequencing

#### Animal harvesting and RNA extraction

Adult hermaphrodites were allowed to lay eggs for six hours. Synchronized progeny were grown on NGM plates seeded with OP50 for ∼30 hrs (L4) at 20°C. For transcriptomic comparisons of PQ treated to untreated control animals, roughly 1200 L4 stage animals were selected for each genotype. Day 1 adult animals were placed on either PQ (4 mM) or control plates the following day. After 48 hours, PQ-treated and control animals were harvested. Worm pellets were snap-frozen in liquid nitrogen and stored at -80°C. Animals were lysed in 10% SDS, b-ME, 0.5 M EDTA, 1M Tris-HCl pH 7.5, 20 mg/ml Proteinase K. Total RNA was extracted with Trizol (ThermoFisher) and purified by RNeasy columns (QIAGEN). RNA samples were frozen using liquid nitrogen and stored at -80°C.

### Differential expression analysis and functional analysis of enriched genes

Quality checks of extracted RNA, cDNA synthesis, and preparation of libraries with fragmenting cDNA were performed by Novogene. Library selection was poly A tail-based, and a paired layout was used to prepare the libraries. mRNA was sequenced using the Illumina platform with Novaseq6000 instrument. FASTQ files were trimmed using Trimmomatic software and mapped onto reference *C. elegans* genome using STAR. All detected transcripts were quantified using RSEM, and differential expression was analyzed using DESeq2. Cutoffs for differential expression were set to be fold change>2, p_adj_<0.01, FDR<0.01. Gene enrichment analysis was performed using WormCat 2.0 (http://www.wormcat.com) (Holdorf et al., 2020). A web-based bioinformatics tool (https://bioinformatics.psb.ugent.be/webtools/Venn/) was used to calculate the overlap of gene sets and to draw Venn diagrams. Venn diagrams were modified using Affinity Designer (Universal-Version 1.0.0). Statistical comparisons of overlapping gene sets were performed by hypergeometric test (http://nemates.org/MA/progs/overlap_stats.html).

### Confocal imaging

Confocal imaging of animals expressing *gar-3::SL2::GFP* was performed using a Yokogawa CSU-X1 spinning disk confocal on a Nikon NiE upright motorized microscope, Nikon 60x oil immersion objective and Prime BSI Express CMOS camera (Teledyne Vision Solutions). Images were acquired using Nikon Elements software. Day 1 adult animals were immobilized with levamisole (2 mM) on 5% agarose pads. For co-localization analyses, the endogenous *gar-3* reporter was crossed into either a transgenic cholinergic (*acr-2p::mCherry*) or GABAergic (*unc-47p::mCherry*) reporter. Whole animal volumes were imaged using the large-image tiling function. Acquisition (488 nm laser, GFP channel): power 5%, exposure time 800 ms, multiplier 25. 561 nm laser (mCherry channel): power 0.5%, exposure time 100 ms, multiplier 10. Imaging of the *nhr-185::GFP* reporter was performed on the same system using a Nikon 4x air objective. Day 3 adult animals exposed to PQ (48 hrs) and age-matched controls were imaged. Acquisition (488 nm, GFP channel): power 20%, exposure time 200 ms. For all measurements, a rectangular ROI of fixed size surrounding the head of the animal was used to quantify total fluorescence from a maximum intensity confocal projection. Confocal imaging of animals expressing *gar-3p::GFPnovo2* was performed using a Yokogawa CSU-10 spinning disk confocal on an Olympus BX51WI microscope equipped with a Hamamatsu C9100-13 EMCCD camera and a 63x oil immersion objective. Day 1 adult animals were imaged and colocalization analyses performed as above. Acquisition (488 nm laser, GFP channel): power 20.5%, exposure time 100 ms, sensitivity 174. For 561nm (mCherry channel): power 20.5%, exposure time 150 ms, sensitivity 174. Post-processing and analysis of all images was performed using Fiji ImageJ software (Schindelin et al., 2012).

### RNA isolation and RT-qPCR

Lysis of Day 3 adult wild-type animals exposed to PQ (48 hr) and age-matched was performed in 0.5% SDS, 5% beta mercapto ethanol (β-ME), 10 mM EDTA, 10 mM tris-HCl (pH 7.4), and proteinase K (0.5 mg/ml) before purification of RNA by TRI-Reagent (Sigma-Aldrich). cDNA was produced with Transcriptor First-strand cDNA kits (Roche), and RT-PCR was performed using Kappa SYBR Green 2X Mastermix. qRT-PCR was performed on an Eppendorf RealPlex. cDNA was standardized to *act-1*. Representative experiments from at least three repetitions are shown.

### Contributions Summary

KB and MMF conceptualized and planned the experiments. Preliminary data supporting some studies were provided by JBR. The experiments and analyses were conducted by KB. Confocal imaging was performed by KB, CM and HR. Protein carbonylation experiments were performed in RPW laboratory in collaboration with KW. AM performed qPCR experiments. DH and AKW aided with differential expression analyses and functional categorization of RNA-seq data. KB performed gene enrichment analyses and comparisons.

## Acknowledgements

We would like to thank members of the Francis lab (Shankar Ramachandran, Samuel Liu) for critical reading of the manuscript. We also thank Shankar Ramachandran, Christopher Lambert, Julia L. Russell, Michael Gorczyca, Lorris Ferrari and Will Joyce for technical assistance. Some nematode strains were provided by the Caenorhabditis Genetics Center which is funded by the NIH National Center for Research Resources.

---

**Figure S1.**
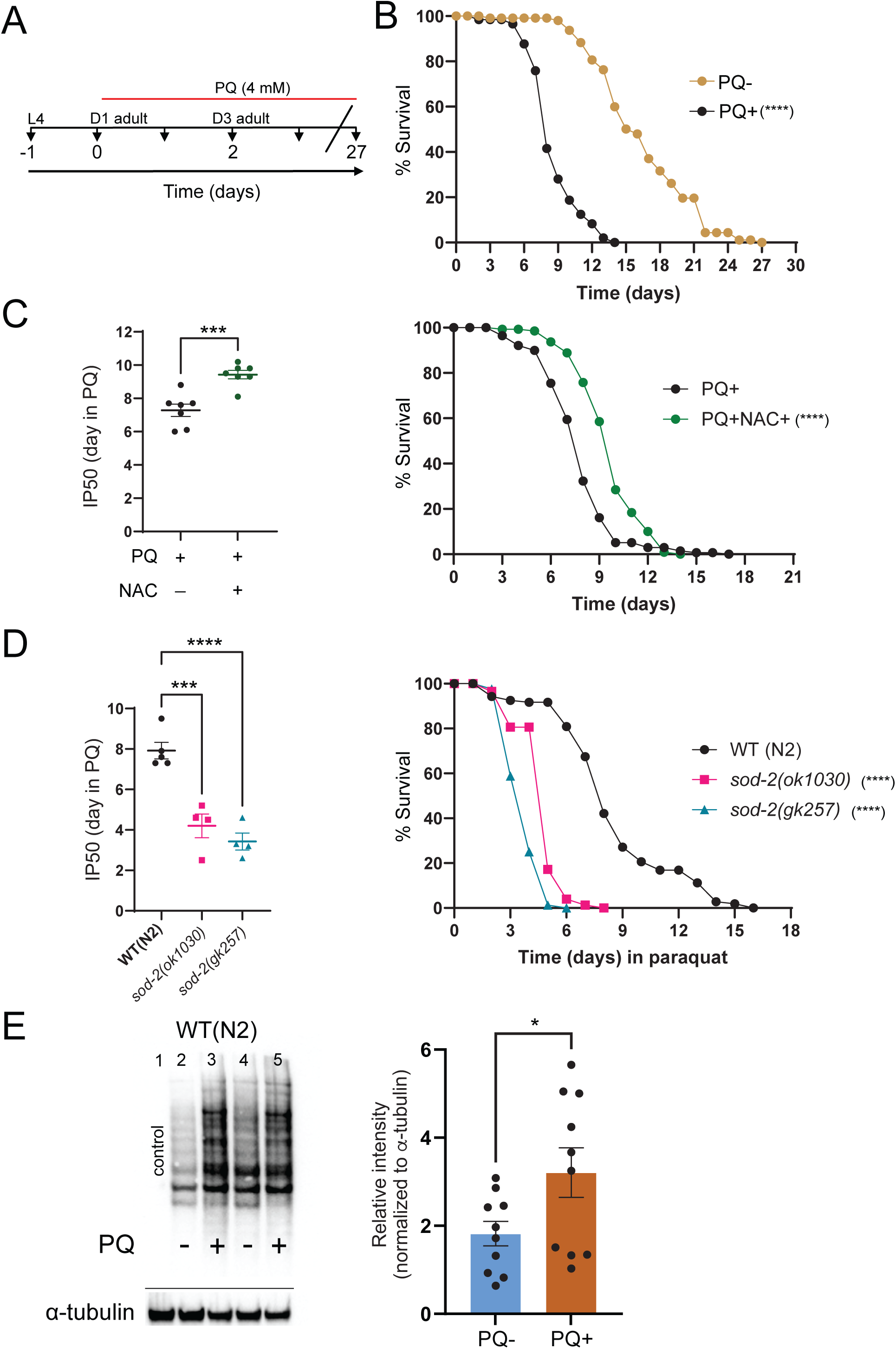
Chronic exposure to paraquat produces oxidative stress **(A)** Experimental timeline for paraquat (PQ) chronic exposure assay. Animals are continuously exposed to 4 mM PQ . At day 0 of the experiment, animals are day 1 (D1) adults. **(B)** Chronic PQ exposure reduces *C. elegans* longevity. Kaplan-Meier curves comparing the longevity of wild type (WT) N2 animals under control conditions (PQ–) to survival on 4 mM PQ. Percentage survival (y-axis) is plotted as a function of time (x-axis). *\*\*\*\*p= 1.4E-12*, Fisher’s exact test at 50% of survival. Total number of independent trials: n=6. The Kaplan-Meier curve is cumulative of all trials. Total number of animals over all trials: n= 112 (PQ-) and 136 (PQ+). **(C)** Left: IP50 comparison between wild-type animals exposed to either PQ (PQ+) or PQ+NAC. For all, IP50 indicates the time to reach half of the original population. Each data point is one independent trial. Bars indicate mean ± SEM. *\*\*\*\*p=0.0007*, unpaired two-tailed t-test with Welch’s correction. Total number of independent trials n=7. Right: Kaplan-Meier survival curves comparing the profiles of animals exposed to either PQ (PQ+) or PQ+NAC . The antioxidant N-acetyl cysteine (NAC) enhances survival during chronic PQ treatment. Kaplan-Meier curve is cumulative of 7 trials. *\*\*\*\*p= 3.60E-12*, Fisher’s exact test at 50% of survival. Total animals: n= 172 (PQ+), n=175 (PQ+NAC). **(D)** Left: IP50 comparison of wild type, *sod-2(ok1030)* and *sod-2(gk257)* treated with PQ. *sod-2* mutants are hypersensitive to PQ. Each data point is one independent trial. Bars indicate mean ± SEM. *\*\*\*p= 0.0004 sod-2(ok1030), ****p<0.0001 sod-2(gk257),* one- way ANOVA with Dunnett’s multiple comparisons test. Total number of trials: n=5 (WT), n=4 for *sod-2(ok1030)* and *sod-2(gk257).* Right: Kaplan-Meier survival curves comparing wild type, *sod-2(ok1030)* and *sod-2(gk257)*. SOD-2 deficiency reduces survival during chronic PQ exposure. Kaplan-Meier curve is cumulative of at least 4 trials. *\*\*\*\*p= 1.1E-12*, *sod-2(ok1030)*, *\*\*\*\*p=4.5E-13*, *sod-2(gk257)*. Fisher’s exact test at 50% of survival. Total number of animals: n= 123, WT; 91, *sod-2(ok1030)*; 90, *sod-2(gk257*). **(E)** Immunoblotting data of protein carbonylation in wild type animals after 48 hr of PQ exposure (starting from D1 adult) compared to untreated controls. Left, Chemiluminescence image of the blot with primary antibody against dinitrophenylhydrazone (DNP) (top) and α-tubulin control (bottom). Right, The ratio of the area under the curve for DNP chemiluminescence in a particular lane relative to α-tubulin. Each data point indicates one trial. Bars indicate mean ± SEM. *\*p=0.0454*, unpaired two-tailed t-test with Welch’s correction.

**Figure S2.**
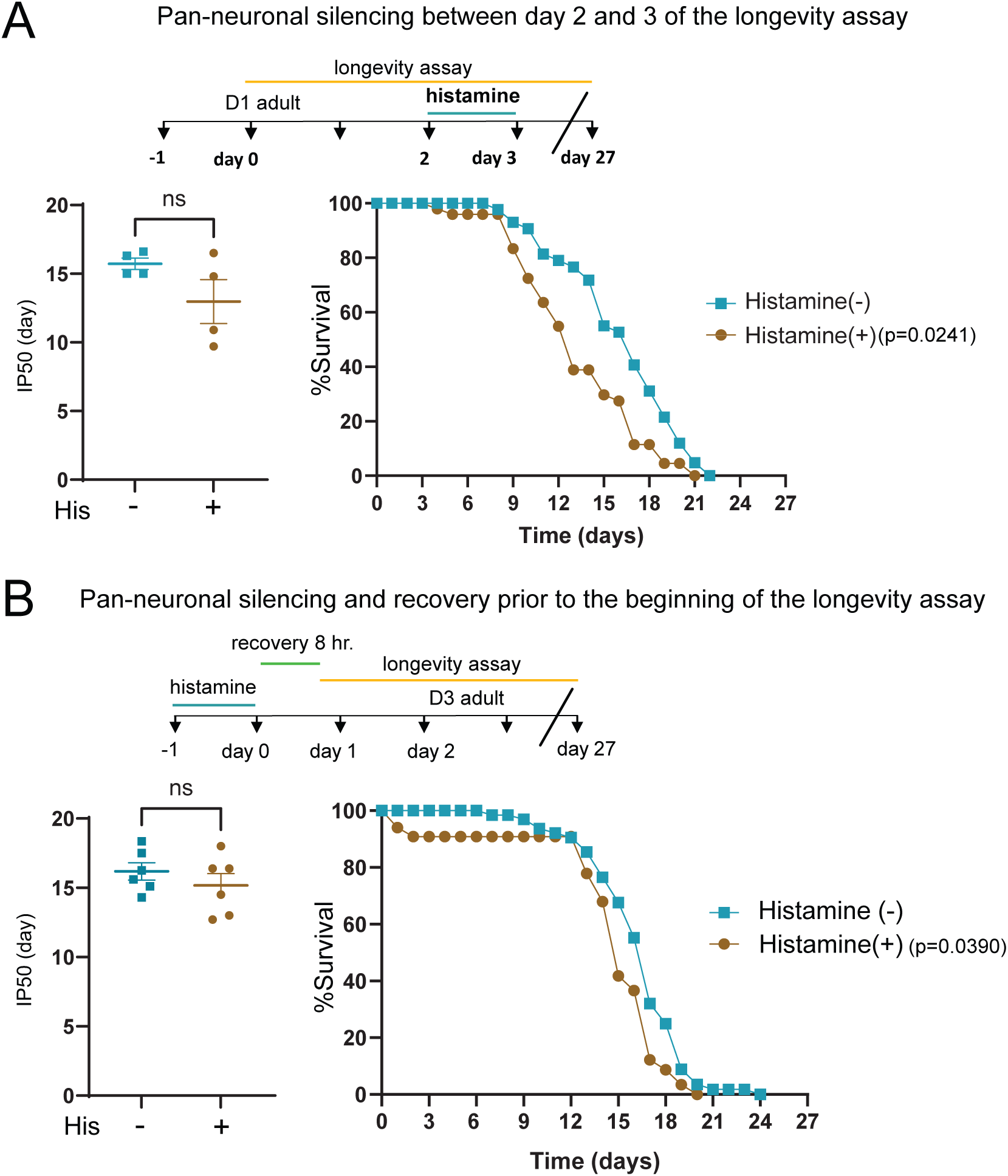
Pan-neuronal silencing of neuronal activity modestly decreases longevity in the absence of PQ exposure **(A)** Top, experimental timeline indicating the period of histamine-mediated neuronal silencing (24 hrs) at day 2 of the longevity assay. Left: Bar graph comparing IP50 measurements for control (His-) and His-treated (His+) *kyEx4571* (*tag-168p*::HisCl1::SL2::GFP) transgenic animals under control conditions (PQ-). The mean IP50 value is not significantly altered by histamine exposure, though variability increases in the His-treated group. Each data point represents one independent trial. Bars indicate mean ± SEM. *p=0.1843 (ns)*, unpaired two-tailed t-test with Welch’s correction. Number of trials: n=4. Right: Kaplan-Meier survival curves for control (His-) and His-treated (His+) groups. Modest differences in longevity are apparent after day 9 of the lifespan assay. Kaplan-Meier curves are cumulative of 4 trials. *\*p = 0.0241*, Fisher’s exact test at 50% of survival. Total number of animals: n= 46 (His-), 49 (His+). **(B)** Top, experimental timeline indicating the periods of histamine-mediated neuronal silencing (24 hr) and recovery (8 hr). Left: Bar graph comparing IP50 measurements for control (His-) and His-treated (His+) *kyEx4571* (*tag-168p*::HisCl1::SL2::GFP) transgenic animals under control conditions (PQ-). Each data point represents one independent trial. Bars indicate mean ± SEM. p=0.3629 (ns), unpaired two-tailed t-test with Welch’s correction. Number of trials, n=6. Right: Kaplan-Meier survival curves comparing profiles of control (His-) and His-treated (His+). Kaplan-Meier curves are cumulative of 6 trials. *\*p = 0.0390*, Fisher’s exact test at 50% of survival. Total number of animals: n=68 (His-),67 (His+).

**Figure S3.**
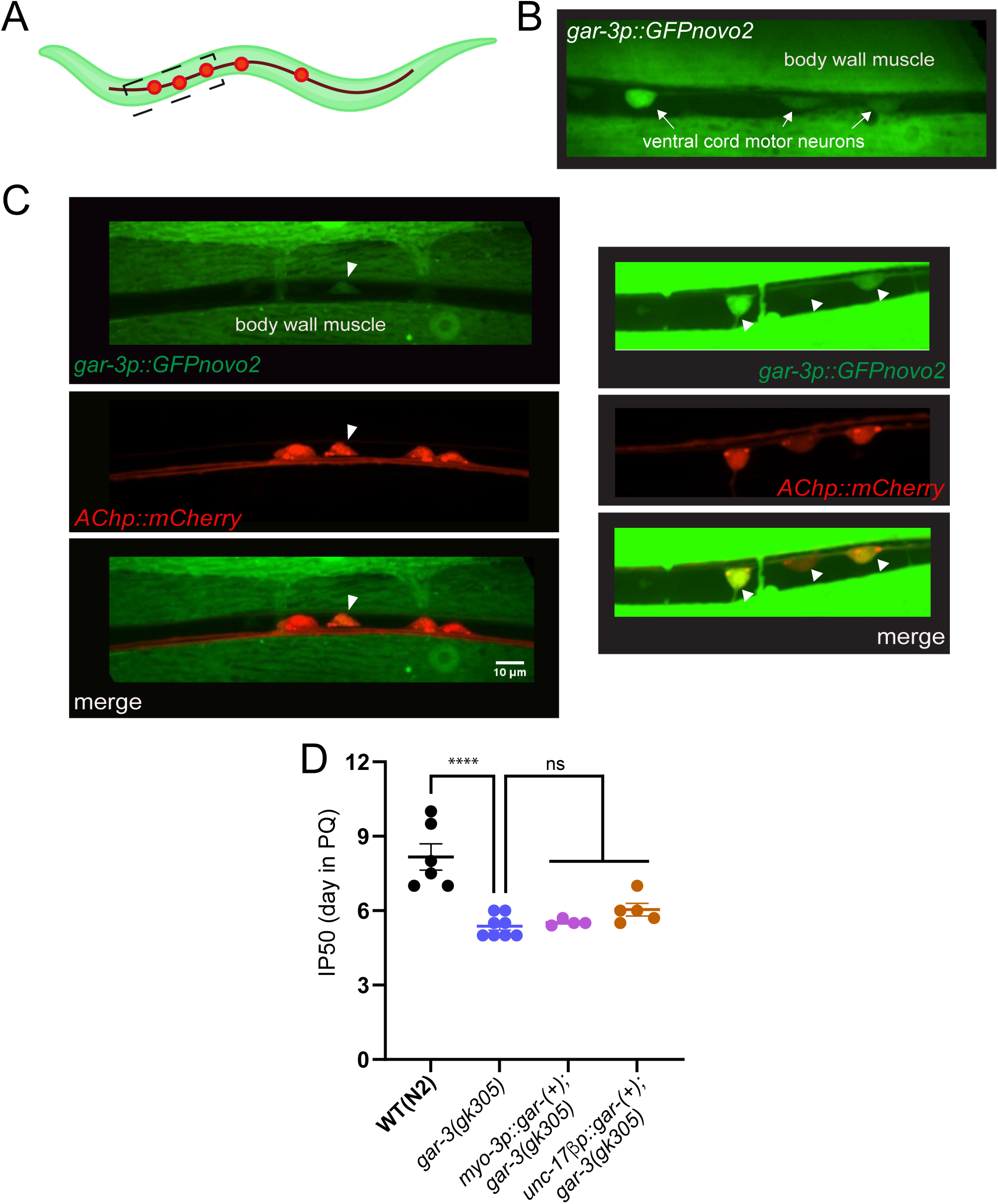
Specific rescue of *gar-3* expression in either cholinergic motor neurons or muscles does not significantly improve IP50 measurements **(A)** Schematic of worm showing area imaged in panels B and C. **(B)** Confocal image showing the expression of a *gar-3p::GFPnovo2* transcriptional reporter in body wall muscles and ventral nerve cord motor neurons. **(C)** Confocal images of ventral nerve cord region showing combined expression of *gar-3p::GFPnovo2* and *Punc-17β::mCherry* reporters. Right image shows increased exposure to better appreciate *gar-3p::GFPnovo2* expression in cholinergic motor neurons. Arrowheads, cholinergic motor neurons expressing *gar-3p::GFPnovo2*. **(D)** Bar graph comparing IP50 measurements for wild type, *gar-3* mutant, ACh motor neuron and muscle-specific rescue strains. Specific expression of wild type *gar-3* cDNA (10 ng/µl) in either ACh motor neurons or muscles of *gar-3* mutants does not offer significant rescue based on IP50 measurement.

**Figure S4.**
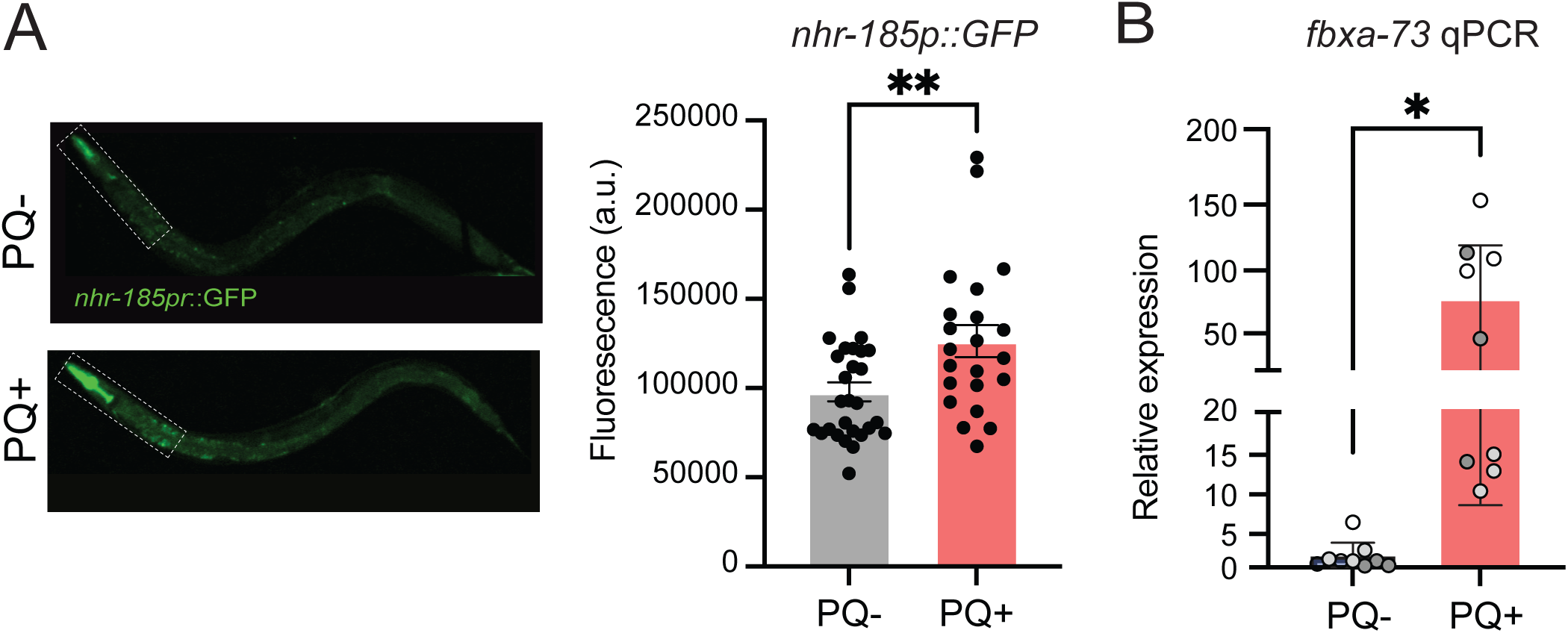
Validation of stress-responsive upregulation of candidate genes identified by RNA-seq. **(A)** Representative images (left) and bar graph (right) quantifying *nhr-185p::GFP* total fluorescence in the pharynx and anterior intestine. Dashed box indicates region of interest for quantification. *nhr-185p::GFP* fluorescence is significantly increased after 48 hrs of PQ (4 mM) treatment. Each data point indicates a measurement from an independent animal. Bars indicate mean ±SEM. *\*\*p<0.01*, Welch’s t-test. PQ-, n=28; PQ+, n=22. **(B)** Quantitative RT-PCR analysis of oxidative stress-responsive *fbxa-73* expression. Prolonged (48 hr) PQ exposure produces a significant increase in *fbxa-73* expression. Each point indicates an independent technical replicate, normalized to *act-1* levels. Bars represent mean ± SEM. *\*p=0.0125*, student’s t-test.

## SUPPLEMENTAL TABLES

Table S1: Differentially expressed genes in wild type (N2) under oxidative stress and their functional categories

Table S2: Classification of stress-related genes upregulated in wild type (N2) under PQ-mediated oxidative stress

Table S3: Comparison of the genes upregulated by arsenite treatment to the genes upregulated by PQ treatment

Table S4: Comparison of the genes upregulated by tBOOH treatment to the genes upregulated by PQ treatment

Table S5: Comparison of the genes upregulated by juglone treatment to the genes upregulated by PQ treatment

Table S6: Differentially expressed genes in unc-17(e113) compared to the wild type (N2) in basal condition (w/o PQ)

Table S7: Differentially expressed genes in *unc-17(e113)* under oxidative stress and their functional categories

Table S8: Overlap of upregulated genes between WT(N2) and *unc-17(e113)*

Table S9: Genes that failed to upregulate in *unc-17(e113)* under PQ-mediated stress are largely different from differentially expressed genes in basal condition (w/o PQ)

Table S10: Differentially expressed genes in *gar-3(gk305)* under PQ-mediated oxidative stress and their functional categories

Table S11: Functional enrichment of genes that fail to upregulate under PQ exposure in either *unc-17(e113)* or *gar-3(gk305)*

Table S12: Genes that are upregulated in all three genotypes N2, *unc-17(e113),* and *gar-3(gk305)* in the presence of PQ

Table S13: Strain list **Table S14:** Plasmid list **Table S15:** Primer list

## REFERENCES

1. Alfonso, A., Grundahl, K., Duerr, J. S., Han, H. P., and Rand, J. B. (1993). The *Caenorhabditis elegans unc-17* gene: a putative vesicular acetylcholine transporter. Science 261, 617–619.

2. Back, P., De Vos, W. H., Depuydt, G. G., Matthijssens, F., Vanfleteren, J. R., and Braeckman, B. P. (2012). Exploring real-time in vivo redox biology of developing and aging Caenorhabditis elegans. Free Radic Biol Med 0, 850–859.

3. Bar-Ziv, R., Dutta, N., Hruby, A., Sukarto, E., Averbukh, M., Alcala, A., Henderson, H. R., Durieux, J., Tronnes, S. U., Ahmad, Q., Bolas, T., Perez, J., Dishart, J. G., Vega, M., Garcia, G., Higuchi-Sanabria, R., and Dillin, A. (2023). Glial-derived mitochondrial signals affect neuronal proteostasis and aging. Sci Adv 9, eadi1411.

4. Bargmann, C. I. (1998). Neurobiology of the Caenorhabditis elegans genome. Science 282, 2028–2033.

5. Bhatti, J. S., Sehrawat, A., Mishra, J., Sidhu, I. S., Navik, U., Khullar, N., Kumar, S., Bhatti, G. K., and Reddy, P. H. (2022). Oxidative stress in the pathophysiology of type 2 diabetes and related complications: Current therapeutics strategies and future perspectives. Free Radic Biol Med 184, 114–134.

6. Biala, Y., Liewald, J. F., Ben-Ami, H. C., Gottschalk, A., and Treinin, M. (2009). The conserved RIC-3 coiled-coil domain mediates receptor-specific interactions with nicotinic acetylcholine receptors. Mol Biol Cell 20, 1419–1427.

7. Biswas, K., Alexander, K., and Francis, M. M. (2022). Reactive Oxygen Species: Angels and Demons in the Life of a Neuron. NeuroSci 3, 130–145.

8. Blackwell, T. K., Steinbaugh, M. J., Hourihan, J. M., Ewald, C. Y., and Isik, M. (2015). SKN-1/Nrf, stress responses, and aging in Caenorhabditis elegans. Free Radic Biol Med 88, 290–301.

9. Cao, X. (2016). Self-regulation and cross-regulation of pattern-recognition receptor signalling in health and disease. Nat Rev Immunol 16, 35–50.

10. Chan, J. P., Hu, Z., and Sieburth, D. (2012). Recruitment of sphingosine kinase to presynaptic terminals by a conserved muscarinic signaling pathway promotes neurotransmitter release. Genes Dev 26, 1070–1085.

11. Chan, J. P., Staab, T. A., Wang, H., Mazzasette, C., Butte, Z., and Sieburth, D. (2013). Extrasynaptic muscarinic acetylcholine receptors on neuronal cell bodies regulate presynaptic function in Caenorhabditis elegans. J Neurosci 33, 14146–14159.

12. Chen, Z.-R., Huang, J.-B., Yang, S.-L., and Hong, F.-F. (2022). Role of Cholinergic Signaling in Alzheimer’s Disease. Molecules 27, 1816.

13. Chun, L., Gong, J., Yuan, F., Zhang, B., Liu, H., Zheng, T., Yu, T., Xu, X. Z., and Liu, J. (2015). Metabotropic GABA signalling modulates longevity in C. elegans. Nat Commun 6, 8828.

14. Cornell, R., Cao, W., Harradine, B., Godini, R., Handley, A., and Pocock, R. (2024). Neuro-intestinal acetylcholine signalling regulates the mitochondrial stress response in Caenorhabditis elegans. Nat Commun 15, 6594.

15. Davies, P., and Maloney, A. J. (1976). Selective loss of central cholinergic neurons in Alzheimer’s disease. Lancet 2, 1403.

16. De Sarno, P., Shestopal, S. A., King, T. D., Zmijewska, A., Song, L., and Jope, R. S. (2003). Muscarinic receptor activation protects cells from apoptotic effects of DNA damage, oxidative stress, and mitochondrial inhibition. J Biol Chem 278, 11086–11093.

17. De Sarno, P., Shestopal, S. A., Zmijewska, A. A., and Jope, R. S. (2005). Anti-apoptotic effects of muscarinic receptor activation are mediated by Rho kinase. Brain Res 1041, 112–115.

18. Desjardins, D., Cacho-Valadez, B., Liu, J. L., Wang, Y., Yee, C., Bernard, K., Khaki, A., Breton, L., and Hekimi, S. (2017). Antioxidants reveal an inverted U-shaped dose-response relationship between reactive oxygen species levels and the rate of aging in Caenorhabditis elegans. Aging Cell 16, 104–112.

19. Doering, K. R. S., Cheng, X., Milburn, L., Ratnappan, R., Ghazi, A., Miller, D. L., and Taubert, S. (2022). Nuclear hormone receptor NHR-49 acts in parallel with HIF-1 to promote hypoxia adaptation in Caenorhabditis elegans. Elife 11, e67911.

20. Dues, D. J., Andrews, E. K., Schaar, C. E., Bergsma, A. L., Senchuk, M. M., and Van Raamsdonk, J. M. (2016). Aging causes decreased resistance to multiple stresses and a failure to activate specific stress response pathways. Aging (Albany NY) 8, 777–795.

21. Eglen, R. M. (2006). Muscarinic receptor subtypes in neuronal and non-neuronal cholinergic function. Auton Autacoid Pharmacol 26, 219–233.

22. Frinchi, M., Nuzzo, D., Scaduto, P., Di Carlo, M., Massenti, M. F., Belluardo, N., and Mudò, G. (2019). Anti-inflammatory and antioxidant effects of muscarinic acetylcholine receptor (mAChR) activation in the rat hippocampus. Sci Rep 9, 14233.

23. Gallrein, C., Williams, A. B., Meyer, D. H., Messling, J.-E., Garcia, A., and Schumacher, B. (2023). baz-2 enhances systemic proteostasis in vivo by regulating acetylcholine metabolism. Cell Rep 42, 113577.

24. Giordano, G., Li, L., White, C. C., Farin, F. M., Wilkerson, H. W., Kavanagh, T. J., and Costa, L. G. (2009). Muscarinic receptors prevent oxidative stress-mediated apoptosis induced by domoic acid in mouse cerebellar granule cells. J Neurochem 109, 525–538.

25. Goh, G. Y. S., Winter, J. J., Bhanshali, F., Doering, K. R. S., Lai, R., Lee, K., Veal, E. A., and Taubert, S. (2018). NHR-49/HNF4 integrates regulation of fatty acid metabolism with a protective transcriptional response to oxidative stress and fasting. Aging Cell 17, e12743.

26. Goswamy, D., and Irazoqui, J. E. (2021). A unifying hypothesis on the central role of reactive oxygen species in bacterial pathogenesis and host defense in C. elegans. Curr Opin Immunol 68, 9–20.

27. Goudeau, J., and Aguilaniu, H. (2010). Carbonylated proteins are eliminated during reproduction in C. elegans. Aging Cell 9, 991–1003.

28. Halevi, S., McKay, J., Palfreyman, M., Yassin, L., Eshel, M., Jorgensen, E., and Treinin, M. (2002). The C. elegans ric-3 gene is required for maturation of nicotinic acetylcholine receptors. EMBO J 21, 1012–1020.

29. Harris, J. J., Jolivet, R., and Attwell, D. (2012). Synaptic energy use and supply. Neuron 75, 762–777.

30. Hipp, M. S., Kasturi, P., and Hartl, F. U. (2019). The proteostasis network and its decline in ageing. Nat Rev Mol Cell Biol 20, 421–435.

31. Holdorf, A. D., Higgins, D. P., Hart, A. C., Boag, P. R., Pazour, G. J., Walhout, A. J. M., and Walker, A. K. (2020). WormCat: An Online Tool for Annotation and Visualization of Caenorhabditis elegans Genome-Scale Data. Genetics 214, 279–294.

32. Hongpaisan, J., Winters, C. A., and Andrews, S. B. (2004). Strong calcium entry activates mitochondrial superoxide generation, upregulating kinase signaling in hippocampal neurons. J Neurosci 24, 10878–10887.

33. Hu, Q., Bian, Q., Rong, D., Wang, L., Song, J., Huang, H.-S., Zeng, J., Mei, J., and Wang, P.-Y. (2023). JAK/STAT pathway: Extracellular signals, diseases, immunity, and therapeutic regimens. Front Bioeng Biotechnol 11, 1110765.

34. Huang, S., Howington, M. B., Dobry, C. J., Evans, C. R., and Leiser, S. F. (2021). Flavin-Containing Monooxygenases Are Conserved Regulators of Stress Resistance and Metabolism. Front Cell Dev Biol 9, 630188.

35. Hwang, J. M., Chang, D. J., Kim, U. S., Lee, Y. S., Park, Y. S., Kaang, B. K., and Cho, N. J. (1999). Cloning and functional characterization of a Caenorhabditis elegans muscarinic acetylcholine receptor. Recept Channels 6, 415–424.

36. Irazoqui, J. E., Troemel, E. R., Feinbaum, R. L., Luhachack, L. G., Cezairliyan, B. O., and Ausubel, F. M. (2010). Distinct pathogenesis and host responses during infection of C. elegans by P. aeruginosa and S. aureus. PLoS Pathog 6, e1000982.

37. Jeong, W.-S., Jun, M., and Kong, A.-N. T. (2006). Nrf2: a potential molecular target for cancer chemoprevention by natural compounds. Antioxid Redox Signal 8, 99–106.

38. Jia, Q., and Sieburth, D. (2021). Mitochondrial hydrogen peroxide positively regulates neuropeptide secretion during diet-induced activation of the oxidative stress response. Nat Commun 12, 2304.

39. Jiang, X., Zhu, D., Okagaki, P., Lipsky, R., Wu, X., Banaudha, K., Mearow, K., Strauss, K. I., and Marini, A. M. (2003). N-methyl-D-aspartate and TrkB receptor activation in cerebellar granule cells: an in vitro model of preconditioning to stimulate intrinsic survival pathways in neurons. Ann N Y Acad Sci 993, 134–45; discussion 159.

40. Juan, C. A., Pérez de la Lastra, J. M., Plou, F. J., and Pérez-Lebeña, E. (2021). The Chemistry of Reactive Oxygen Species (ROS) Revisited: Outlining Their Role in Biological Macromolecules (DNA, Lipids and Proteins) and Induced Pathologies. Int J Mol Sci 22, 4642.

41. Kahn, N. W., Rea, S. L., Moyle, S., Kell, A., and Johnson, T. E. (2008). Proteasomal dysfunction activates the transcription factor SKN-1 and produces a selective oxidative-stress response in Caenorhabditis elegans. Biochem J 409, 205–213.

42. Kato, M., Kolotuev, I., Cunha, A., Gharib, S., and Sternberg, P. W. (2021). Extrasynaptic acetylcholine signaling through a muscarinic receptor regulates cell migration. Proc Natl Acad Sci U S A 118, e1904338118.

43. Koga, H., Kaushik, S., and Cuervo, A. M. (2011). Protein homeostasis and aging: The importance of exquisite quality control. Ageing Res Rev 10, 205–215.

44. Kratsios, P., Stolfi, A., Levine, M., and Hobert, O. (2011). Coordinated regulation of cholinergic motor neuron traits through a conserved terminal selector gene. Nat Neurosci 15, 205–214.

45. Labed, S. A., Wani, K. A., Jagadeesan, S., Hakkim, A., Najibi, M., and Irazoqui, J. E. (2018). Intestinal Epithelial Wnt Signaling Mediates Acetylcholine-Triggered Host Defense against Infection. Immunity 48, 963–978.e3.

46. Leiser, S. F., Miller, H., Rossner, R., Fletcher, M., Leonard, A., Primitivo, M., Rintala, N., Ramos, F. J., Miller, D. L., and Kaeberlein, M. (2015). Cell nonautonomous activation of flavin-containing monooxygenase promotes longevity and health span. Science 350, 1375–1378.

47. Liu, J., Narasimhan, P., Yu, F., and Chan, P. H. (2005). Neuroprotection by hypoxic preconditioning involves oxidative stress-mediated expression of hypoxia-inducible factor and erythropoietin. Stroke 36, 1264–1269.

48. Liu, Y., Zhou, J., Zhang, N., Wu, X., Zhang, Q., Zhang, W., Li, X., and Tian, Y. (2022). Two sensory neurons coordinate the systemic mitochondrial stress response via GPCR signaling in C. elegans. Dev Cell 57, 2469–2482.e5.

49. Ma, T., Hoeffer, C. A., Wong, H., Massaad, C. A., Zhou, P., Iadecola, C., Murphy, M. P., Pautler, R. G., and Klann, E. (2011). Amyloid β-induced impairments in hippocampal synaptic plasticity are rescued by decreasing mitochondrial superoxide. J Neurosci 31, 5589–5595.

50. Manoharan, S., Guillemin, G. J., Abiramasundari, R. S., Essa, M. M., Akbar, M., and Akbar, M. D. (2016). The Role of Reactive Oxygen Species in the Pathogenesis of Alzheimer’s Disease, Parkinson’s Disease, and Huntington’s Disease: A Mini Review. Oxid Med Cell Longev 2016, 8590578.

51. Maremonti, E., Eide, D. M., Oughton, D. H., Salbu, B., Grammes, F., Kassaye, Y. A., Guédon, R., Lecomte-Pradines, C., and Brede, D. A. (2019). Gamma radiation induces life stage-dependent reprotoxicity in Caenorhabditis elegans via impairment of spermatogenesis. Sci Total Environ 695, 133835.

52. Mattson, M. P., and Cheng, A. (2006). Neurohormetic phytochemicals: Low-dose toxins that induce adaptive neuronal stress responses. Trends Neurosci 29, 632–639.

53. Mello, C. C., Kramer, J. M., Stinchcomb, D., and Ambros, V. (1991). Efficient gene transfer in *C.elegans*: extrachromosomal maintenance and integration of transforming sequences. EMBO J 10, 3959–3970.

54. Mesbahi, H., Pho, K. B., Tench, A. J., Leon Guerrero, V. L., and MacNeil, L. T. (2020). Cuticle Collagen Expression Is Regulated in Response to Environmental Stimuli by the GATA Transcription Factor ELT-3 in Caenorhabditis elegans. Genetics 215, 483–495.

55. Mufson, E. J., Counts, S. E., Perez, S. E., and Ginsberg, S. D. (2008). Cholinergic system during the progression of Alzheimer’s disease: therapeutic implications. Expert Rev Neurother 8, 1703–1718.

56. Murphy, C. T., McCarroll, S. A., Bargmann, C. I., Fraser, A., Kamath, R. S., Ahringer, J., Li, H., and Kenyon, C. (2003). Genes that act downstream of DAF-16 to influence the lifespan of Caenorhabditis elegans. Nature 424, 277–283.

57. Nandi, I., and Aroeti, B. (2023). Mitogen-Activated Protein Kinases (MAPKs) and Enteric Bacterial Pathogens: A Complex Interplay. Int J Mol Sci 24, 11905.

58. Nasrallah, M. A., Peterson, N. D., Szumel, E. S., Liu, P., Page, A. L., Tse, S. Y., Wani, K. A., Tocheny, C. E., and Pukkila-Worley, R. (2023). Transcriptional suppression of sphingolipid catabolism controls pathogen resistance in C. elegans. PLoS Pathog 19, e1011730.

59. Nguyen, T., Nioi, P., and Pickett, C. B. (2009). The Nrf2-antioxidant response element signaling pathway and its activation by oxidative stress. J Biol Chem 284, 13291–13295.

60. Oliveira, R. P., Porter Abate, J., Dilks, K., Landis, J., Ashraf, J., Murphy, C. T., and Blackwell, T. K. (2009). Condition-adapted stress and longevity gene regulation by Caenorhabditis elegans SKN-1/Nrf. Aging Cell 8, 524–541.

61. Papaevgeniou, N., and Chondrogianni, N. (2014). The ubiquitin proteasome system in Caenorhabditis elegans and its regulation. Redox Biol 2, 333–347.

62. Paulsen, C. E., Truong, T. H., Garcia, F. J., Homann, A., Gupta, V., Leonard, S. E., and Carroll, K. S. (2011). Peroxide-dependent sulfenylation of the EGFR catalytic site enhances kinase activity. Nat Chem Biol 8, 57–64.

63. Petrash, H. A., Philbrook, A., Haburcak, M., Barbagallo, B., and Francis, M. M. (2013). ACR-12 ionotropic acetylcholine receptor complexes regulate inhibitory motor neuron activity in *Caenorhabditis elegans*. J Neurosci 33, 5524–5532.

64. Pokala, N., Liu, Q., Gordus, A., and Bargmann, C. I. (2014). Inducible and titratable silencing of *Caenorhabditis elegans* neurons in vivo with histamine-gated chloride channels. Proc Natl Acad Sci U S A 111, 2770–2775.

65. Robertson, R. M. (2004). Modulation of neural circuit operation by prior environmental stress. Integr Comp Biol 44, 21–27.

66. Sandhu, A., Badal, D., Sheokand, R., Tyagi, S., and Singh, V. (2021). Specific collagens maintain the cuticle permeability barrier in Caenorhabditis elegans. Genetics 217, iyaa047.

67. Schaar, C. E., Dues, D. J., Spielbauer, K. K., Machiela, E., Cooper, J. F., Senchuk, M., Hekimi, S., and Van Raamsdonk, J. M. (2015). Mitochondrial and cytoplasmic ROS have opposing effects on lifespan. PLoS Genet 11, e1004972.

68. Schindelin, J., Arganda-Carreras, I., Frise, E., Kaynig, V., Longair, M., Pietzsch, T., Preibisch, S., Rueden, C., Saalfeld, S., Schmid, B., Tinevez, J.-Y., White, D. J., Hartenstein, V., Eliceiri, K., Tomancak, P., and Cardona, A. (2012). Fiji: an open-source platform for biological-image analysis. Nat Methods 9, 676–682.

69. Senchuk, M. M., Dues, D. J., and Van Raamsdonk, J. M. (2017). Measuring Oxidative Stress in Caenorhabditis elegans: Paraquat and Juglone Sensitivity Assays. Bio Protoc 7, e2086.

70. Serrano-Saiz, E., Gulez, B., Pereira, L., Gendrel, M., Kerk, S. Y., Vidal, B., Feng, W., Wang, C., Kratsios, P., Rand, J. B., and Hobert, O. (2020). Modular Organization of Cis-regulatory Control Information of Neurotransmitter Pathway Genes in Caenorhabditis elegans. Genetics 215, 665–681.

71. Silva, M. C., Amaral, M. D., and Morimoto, R. I. (2013). Neuronal reprograming of protein homeostasis by calcium-dependent regulation of the heat shock response. PLoS Genet 9, e1003711.

72. Smith, J. J., Taylor, S. R., Blum, J. A., Feng, W., Collings, R., Gitler, A. D., Miller, D. M., and Kratsios, P. (2024). A molecular atlas of adult C. elegans motor neurons reveals ancient diversity delineated by conserved transcription factor codes. Cell Rep 43, 113857.

73. Steger, K. A., and Avery, L. (2004). The GAR-3 muscarinic receptor cooperates with calcium signals to regulate muscle contraction in the Caenorhabditis elegans pharynx. Genetics 167, 633–643.

74. Tanaka, K. (2009). The proteasome: overview of structure and functions. Proc Jpn Acad Ser B Phys Biol Sci 85, 12–36.

75. Taylor, S. R., Santpere, G., Weinreb, A., Barrett, A., Reilly, M. B., Xu, C., Varol, E., Oikonomou, P., Glenwinkel, L., McWhirter, R., Poff, A., Basavaraju, M., Rafi, I., Yemini, E., Cook, S. J., Abrams, A., Vidal, B., Cros, C., Tavazoie, S., Sestan, N., Hammarlund, M., Hobert, O., and Miller, D. M. (2021). Molecular topography of an entire nervous system. Cell 184, 4329–4347.e23.

76. Tullet, J. M., Hertweck, M., An, J. H., Baker, J., Hwang, J. Y., Liu, S., Oliveira, R. P., Baumeister, R., and Blackwell, T. K. (2008). Direct inhibition of the longevity-promoting factor SKN-1 by insulin-like signaling in C. elegans. Cell 132, 1025–1038.

77. Valko, M., Izakovic, M., Mazur, M., Rhodes, C. J., and Telser, J. (2004). Role of oxygen radicals in DNA damage and cancer incidence. Mol Cell Biochem 266, 37–56.

78. van Heemst, D. (2010). Insulin, IGF-1 and longevity. Aging Dis 1, 147–157.

79. Van Raamsdonk, J. M., and Hekimi, S. (2009). Deletion of the mitochondrial superoxide dismutase sod-2 extends lifespan in Caenorhabditis elegans. PLoS Genet 5, e1000361.

80. Van Raamsdonk, J. M., and Hekimi, S. (2012). Superoxide dismutase is dispensable for normal animal lifespan. Proc Natl Acad Sci U S A 109, 5785–5790.

81. Wani, K. A., Goswamy, D., and Irazoqui, J. E. (2020). Nervous system control of intestinal host defense in C. elegans. Curr Opin Neurobiol 62, 1–9.

82. Wani, K. A., Goswamy, D., Taubert, S., Ratnappan, R., Ghazi, A., and Irazoqui, J. E. (2021). NHR-49/PPAR-α and HLH-30/TFEB cooperate for C. elegans host defense via a flavin-containing monooxygenase. Elife 10, e62775.

83. West, A. E., Chen, W. G., Dalva, M. B., Dolmetsch, R. E., Kornhauser, J. M., Shaywitz, A. J., Takasu, M. A., Tao, X., and Greenberg, M. E. (2001). Calcium regulation of neuronal gene expression. Proc Natl Acad Sci U S A 98, 11024–11031.

84. Wu, C.-W., Deonarine, A., Przybysz, A., Strange, K., and Choe, K. P. (2016). The Skp1 Homologs SKR-1/2 Are Required for the Caenorhabditis elegans SKN-1 Antioxidant/Detoxification Response Independently of p38 MAPK. PLoS Genet 12, e1006361.

85. Yang, J.-S., Nam, H.-J., Seo, M., Han, S. K., Choi, Y., Nam, H. G., Lee, S.-J., and Kim, S. (2011). OASIS: online application for the survival analysis of lifespan assays performed in aging research. PLoS One 6, e23525.

86. Yang, W. S., Kim, K. J., Gaschler, M. M., Patel, M., Shchepinov, M. S., and Stockwell, B. R. (2016). Peroxidation of polyunsaturated fatty acids by lipoxygenases drives ferroptosis. Proc Natl Acad Sci U S A 113, E4966–75.

87. Zhang, Y., Gan, B., Liu, D., and Paik, J.-h. (2011). FoxO family members in cancer. Cancer Biol Ther 12, 253–259.

